# The interplay between detection and localization in human vision

**DOI:** 10.64898/2026.07.06.736811

**Authors:** Fabian Coupette, David H. Brainard, Hannah E. Smithson, Daniel J. Read

## Abstract

Detection and localization are fundamental but distinct tasks of sensory systems: detecting a signal requires accumulating evidence for its presence, whereas localizing it requires extracting spatial information. How active sampling should be organized when both tasks must be performed simultaneously remains unclear. Here, using an analytically tractable model of early visual processing and Bayesian ideal-observer inference, we establish a direct relation between the two tasks: localizing a stimulus is equivalent to detecting its spatial gradient. Consequently, detection and localization favor fundamentally different sampling dynamics when the stimulus size exceeds the effective blur scale of the visual system. Applied to fixational eye movements (FEMs), which continually translate stationary visual stimuli across the adapting retina, this distinction yields two competing optimal movement scales. Localization is optimized when the eye moves approximately one retinal blur length during the adaptation time, whereas detection is optimized by motion on the scale of the stimulus size. The scale of physiological FEMs lies near the predicted localization optimum, while stimulus detection remains comparatively robust to variations in eye motion. Our model recovers established laws of temporal and spatial summation while predicting additional scaling regimes arising from FEMs. We propose an experimental paradigm that tests these predictions by measuring localization at matched detectability, eliminating the unknown internal-noise amplitude. Our results reveal a fundamental trade-off between acquiring evidence for the presence and position of spatially extended signals and suggest that human fixational eye movements preferentially support spatial localization.

## I. INTRODUCTION

Two of the most essential tasks of the human visual system are detection and localization of stimuli. In common experience, a detected stimulus is perceived to be at a specific location and localization is certainly facilitated by the awareness of something to localize. However, localization conceptually does not necessitate conscious detection as illustrated by studies on blindsight [1–3]. Likewise, detection of a stimulus is notably not always accompanied by an accurate localization, for instance, in the visual periphery [4, 5]. Thus, although the two tasks rarely appear in isolation the two processes are conceptually distinct. The visual system needs to address both tasks but how exactly both functionalities can be addressed efficiently at the same time remains enigmatic.

The human eye is not completely stationary during periods of fixation. This insight dates back to the nineteenth century [6]. However, due to the small scale of fixational eye movements (FEMs), their systematic exploration only began in the 1950s [7, 8]. Contemporary techniques such as the Dual-Purkinje Eye Tracker [9, 10] or the Adaptive Optics Scanning Laser Ophthalmoscope [11–13] are able to track motion on the scale of individual foveal cone photoreceptors. Nonetheless, many fundamental questions regarding the properties and purposes of fixational eye movements remain.

FEMs comprise sudden jumps of up to a degree of visual angle called *microsaccades* as well as a smaller-scale diffusion-like motion between microsaccades referred to as *ocular drift* [8, 14–16]. Here we focus on information acquisition during drift, as microsaccades are relatively infrequent and the eye therefore spends most of the fixation period drifting. Whether fixational drift provides a functional benefit to vision, and what that benefit might be, is the subject of ongoing debate. It has been suggested that FEMs are a mechanical consequence of statistical fluctuations in the oculomotor control system [17, 18] or that they serve to adjust the fixation target to the most acute part of the retina [14, 19]. However, many studies argue that FEMs play a more elaborate role in the processing of visual information [10, 14, 15, 20]. As part of coding the visual information efficiently, neurons adapt to constant light stimulation [16, 21–23], i.e. their sensitivity and responsiveness towards continued stimulation gradually decreases.

Fixational eye movements assist in overcoming the effects of adaptation [16, 24]. As such FEMs may serve as a sampling strategy matched to the specific way that the acquired information is being processed. However, these eye movements also cause the information of a stationary stimulus to be spread out over multiple receptors, decreasing the signal-to-noise ratio of any individual photoreceptor [25–27]. That means, motion that is too rapid may result in insufficient stimulation of individual photoreceptors, while motion that is too slow leads to signal loss due to adaptation. That begs the question: what eye movements optimally supports a specific task of the visual system?

Within this manuscript we first examine the interplay between detection and localization in general before explicitly studying the impact of random eye movements on either operation. To this end, we study a simple analytic model of the retina described in Section II. Using Bayesian inference, we derive the key quantities that characterize detection capability and localization accuracy of an ideal observer in Section III. Finally, we present how these key quantities vary with fixational eye movements in Sec. IV. Section V concludes.

## II. BASELINE MODEL FOR RETINAL RESPONSE

We employ a simplified model of retinal response by Bialek [27, 28]. We model the retina as a continuum of receptors embedded in the real plane R^2^. A stationary stimulus *s*(***r***) is projected onto the retina but subject to blur induced directly by the eye’s optics or indirectly through the finite extent of receptors on the retina or subsequent post-receptoral units. As a consequence, the stimulation that reaches the eye is the convolution of the original stimulus with a blurring function *F*. The movement of the retina ***R***(*t*) effectively translates into a motion of the stimulus relative to the retina so that the instantaneous stimulation of the retina is given by

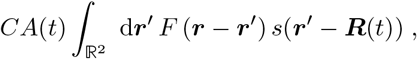

where we include a stimulus modulation via amplitude *A*(*t*) and an overall contrast *C* representing the stimulus intensity relative to a uniformly illuminated background. We extract *C* as a single overall scaling parameter implying that the amplitude function *A* and the function describing the stimulus *s* are reasonably normalized. Within this article we specifically study

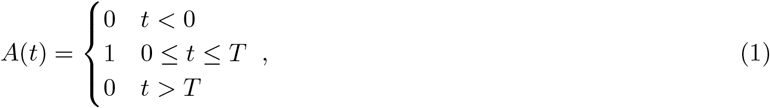

i.e. a finite window of stimulus presentation of duration *T*.

A key feature of retinal response is adaptation to persistent stimulation [29, 30]. To incorporate this, we employ a simple linear model, such that the retinal response depends on all previous stimulations weighted by an adaptation kernel *K*. Causality requires *K*(*t <* 0) = 0, and we additionally demand

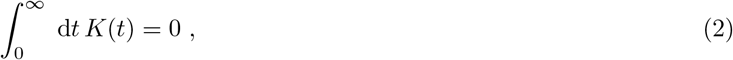

which corresponds to *perfect adaptation*, i.e. a stationary image fades entirely for long observation times.

So far all ingredients of the model are deterministic – knowing the eye movement means we know the receptor response exactly. Yet, each receptor is subject to various kinds of noise limiting the transfer of information. Within this article we model the neuronal fluctuations as Gaussian white noise added to the overall receptor response. There are other sources of stochasticity such as photon shot noise, which becomes a significant factor at low light levels [28, 31–33], or fluctuations in the early stages of photoreceptor excitation. Such noise would be passed through some of the temporal and spatial filtering circuitry represented by the functions *K* and *F*, resulting in colored noise in the final signal. We neglect these sources of colored noise for simplicity. Consequently, the equation for the retinal response reads

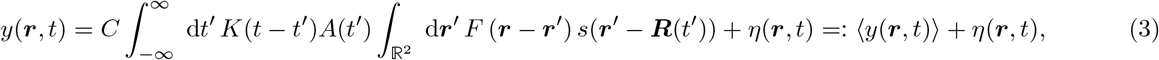

with the noise entirely characterized by the variance *σ*^2^

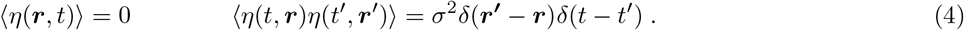

The angular brackets ⟨·⟩ denote the average over all noise realizations corresponding to that variance. The progressive build-up of eq. (3) and the effect of its constituents on the final signal are illustrated in Fig. 1.

**Figure 1.**
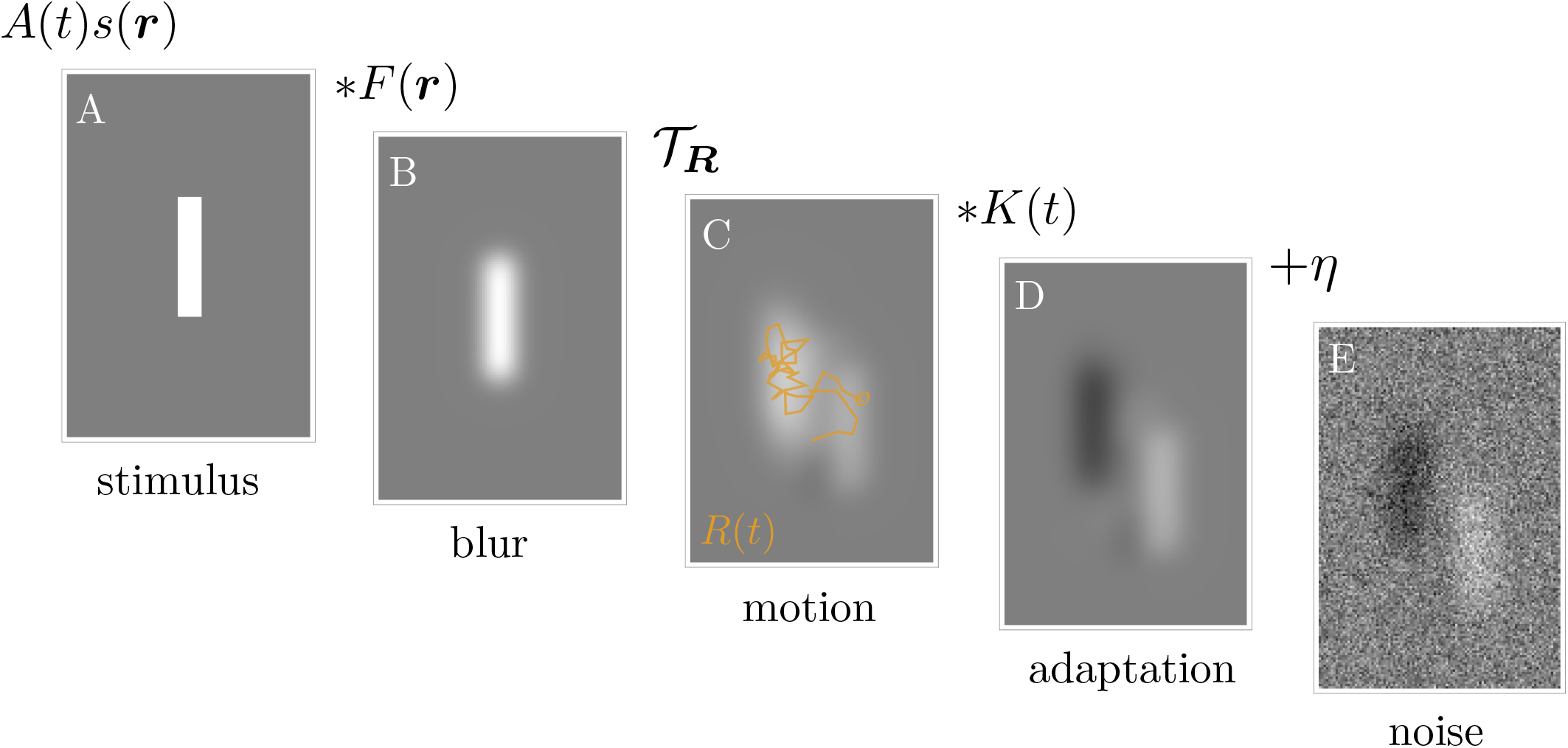
Progressive construction of the baseline model showing retinal excitation after time *T* : A rectangular stimulus (A) is blurred by the eye’s optics to produce the instantaneous retinal image (B). The asterisk denotes a convolution. The blurred stimulus is moved across the retina due to FEMs. *T*_***R***_ represents the translation according to the trajectory ***R***. (C shows the average stimulation between time *t* = 0 and *t* = *T*). Including neural adaptation through a temporal filter *K*, previous stimulation induces a (black) after-image (D). Finally, Gaussian white noise is added (E).

The spatial integral in eq. (3) is a convolution which turns into a simpler product when transforming into spatial frequencies

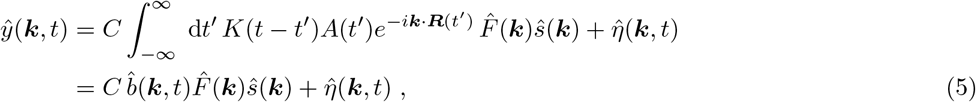

where 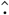 denotes the Fourier transform of the corresponding functions, using the convention 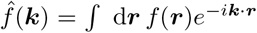. The function

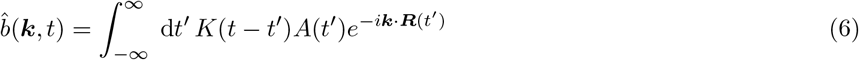

encodes the temporal component of the average receptor response at wavevector ***k***, thus combining the effects of adaptation, stimulus modulation and eye movement.

Concrete calculations require a specification of the functional forms for the temporal filter *K*, and the blurring function *F*. We assume the temporal filter *K* is a band-pass filter, excitatory (positive) at small times and inhibitory (negative) for long times. We also demand perfect adaptation 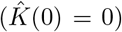. The qualitative behavior of the model is largely insensitive to the exact choice of band pass filter [27] so that we pick a bi-exponential *K* for analytic simplicity. More elaborate expressions based on physiological considerations can be found in [29]. The temporal filter sets a characteristic adaptation time *t*_*a*_ describing the transition from excitatory to inhibitory response to continued stimulation, i.e. a typical timescale over which an image fixed on the retina is integrated positively. We specifically target early visual processing for which *t*_*a*_ ≈ 40 ms is a reasonable estimate at mid photopic light levels [29, 34, 35]. We assume the spatial filter *F* is a low-pass filter, which blurs the incoming signal over a typical length scale *β*. The eye’s optics typically induces blur over lengths of roughly 60″ (arc seconds). However, adaptive optics may account rendered dimensionless by rescaling with the appropriate scales such as for individual aberrations so that the extent of a single cone in the fovea, about 30″ in diameter, becomes the limiting factor. Thus, *β* ≈ 30″ − 60″ or around 1 *µ*m on the retinal surface [36] is a natural range for the blur length. The timescale *t*_*a*_ and the length scale *β* are intrinsic properties of the visual system and may vary across individuals and viewing conditions. Thus, throughout this manuscript we will measure time in units of *t*_*a*_ and length in units of *β*. Accordingly, the variance of the added white noise which is the final intrinsic parameter of our model retina is measured in units of *t*_*a*_*β*^2^. We use the notation 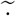 to indicate quantities expressed in **natural units**, i.e. parameters

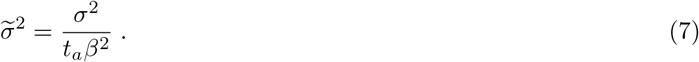

Having set up our model visual system, we proceed by characterizing two key functionalities: detection and localization.

## III. DETECTION AND LOCALIZATION

The most elementary information that can be conveyed about any stimulus is its existence, i.e. whether the stimulus is present or not. Everyday experience suggests that before a stimulus can be characterized, there needs to be some degree of confidence that there indeed is a stimulus. The binary assessment of evidence to decide between stimulus present or absent is referred to as a *detection task* [37]. To remove criterion effects, measurements of detection performance are typically made in *two-alternative-forced-choice* (2AFC) in which the stimulus is presented in one of two intervals or places and the observer must decide which of the two contained the stimulus. The fraction of correct responses in a 2AFC detection task varies with the stimulus contrast giving rise to a *psychometric function* [38] linking an observer’s decisions to the physical parameters of the experiment such as the contrast of the stimulus. In this context, the psychometric function describes the transition between invisibility and visibility of the stimulus and typically has a sigmoidal shape. The critical contrast *C*_crit_ for which the psychometric function exceeds a prescribed value marks the *detection threshold* which can be inferred from experimental measurements of the psychometric function. We may also ask for specific properties of the stimulus such as its position relative to other stimuli in the visual field. This would be a type of *localization task*. The accuracy of localization can, for instance, be probed through a *Vernier task*, a binary judgment of the direction a relative offset between two stimuli [39]. In the same way as for contrast in a detection task, this can be framed as a 2AFC task giving rise to a psychometric function and a critical offset known as the *Vernier* threshold that is a measure of visual localization. In the following, we derive the relationship between these thresholds in the framework of our baseline model of visual perception.

In particular, we study an *ideal observer* (IO) who knows the spatial profile of the set of presented stimuli and their own visual system [40]. In our formulation here this includes the IO’s eye movement trajectory on each trial. For this reason, with the exception of the added noise, the IO can perfectly anticipate the time-dependent retinal excitation of any admissible stimulus. Knowledge about the stimulus is realistic in a detection experiment, yet perfect knowledge of the observer’s visual system and optimal processing is certainly a simplification. We describe the best performance possible given the available information, so the results of our calculations provide lower bounds for threshold measurements in actual observers. Since our model does not distinguish between movement of the eye and movement of the entire visual scene, this trend also applies for moving stimuli. However, that analogy would require the ideal observer to perfectly anticipate the trajectory of the stimulus. Moreover, macroscopic movement is detected and processed through specialized pathways. Thus, awareness of the implicit assumptions is required when treating stimulus motion as inverse eye motion and vice versa.

### III.1. Detection

The IO knows that they are observing one of two conditions: Either the stimulus *s* is present (on), resulting in the noisy spatio-temporal excitation pattern *y*(***r***, *t*) ≡ *y*_on_(***r***, *t*) + *η*(***r***, *t*), or the stimulus is absent (off), resulting in noise alone *y*(***r***, *t*) = *y*_off_ (***r***, *t*) + *η*(***r***, *t*) with *y*_off_ ≡ 0. The only stochastic component is the added Gaussian white noise.

The task of the IO is to infer whether the stimulus is more likely to be “on” or “off”, i.e. the stimulus condition, given the observed signals *y*(***r***, *t*). Hence, under the IO-assumptions we cast the detection task in terms of Bayesian inference. Using the noise statistics, it is straightforward to calculate the probability distribution of the signal *y*(***r***, *t*) given the stimulus condition, “on” or “off”. Bayes’ theorem allows us to reverse this, i.e. to express the posterior probability of each stimulus condition given the set of signals of all receptors at all times *y*(***r***, *t*). We assume the IO infers the stimulus condition depending on which is the more likely, in this Bayesian sense including prior probabilities of either condition. This decision rule maximizes the fraction of correct assessments in the 2AFC task. The derivation is given in Appendix A.

The IO makes the correct decision if they decide “on” if the stimulus was in fact present, or, conversely, decide “off” if the stimulus was absent. If both conditions are occurring with the same prior probability, the probability of a correct decision is:

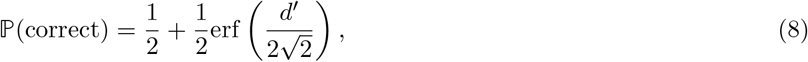

where *d*′ quantifies the difference between the mean excitation patterns generated by the on and off conditions relative to the noise:

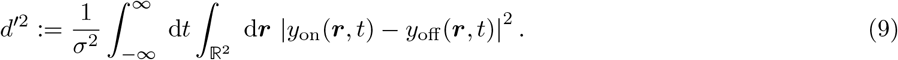

Thus, the simple scalar “signal-to-noise ratio” *d*′ determines the detection capability of the IO. Notably, eq. (9) is invariant under translation, i.e. it does not matter where the stimulus lands on our continuous model retina. Moreover, the IO can never perform worse than guessing, i.e. 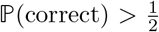. We choose the symbol *d*′ in analogy to classical signal detection theory [41] where it, in the same way as here, characterizes the difference of means of two Gaussian distributions divided by the noise standard deviation. Since *y*_off_ ≡ 0, *d*′ scales linearly with the ratio 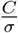. This is an immediate consequence of the linearity of the model (cf. eq. (9)). Thus, we may define

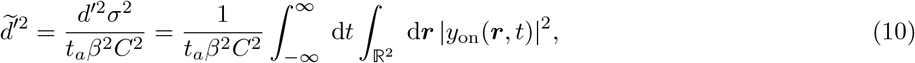

where we scaled out the trivial dependencies and made the result dimensionless by also expressing the variance *σ*^2^ in natural units 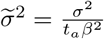. Importantly, 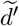 is independent of contrast and noise. Thus, it characterizes the detectability of a stimulus exclusively on the basis of the interaction between stimulus properties, spatial and temporal filters, and fixational eye movements.

Using the model and notation developed in section II and the Plancherel theorem, factoring out the power spectrum of the stationary stimulus yields

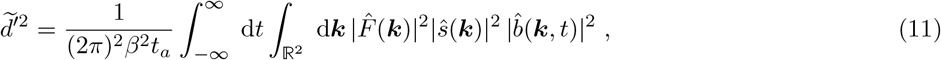

where 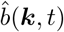 encodes the coupling between eye movements and temporal filter, and 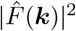 and 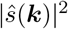 denote the power transfer function of the spatial filter and the power spectrum of the stimulus, respectively. The time-integral of 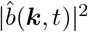 has previously been analyzed as an ensemble average over diffusive trajectories [27] (where it is called *χ*_***k***_), which allows us to simplify eq. (11) to a single integral over ***k***. As this ensemble average is isotropic, we further simplify eq. (11) for radially symmetric stimuli

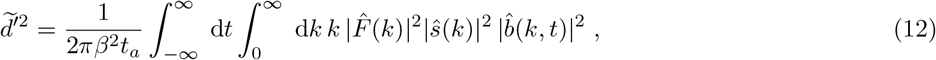

with *k* := |***k***|.

#### Contrast Threshold for Stimulus Detection

When evaluating stimulus detection, a common procedure is to vary the stimulus contrast *C* and determine the threshold *C*_crit_ at which the probability of correct response in a 2AFC detection task exceeds a critical value. In signal detection theory, this occurs when *d*′ attains a critical value *z*_crit_. Hence, we may express the detection threshold as

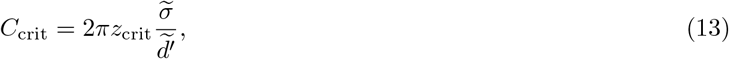

with 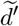 evaluated by eq. (11). While the extent of receptor noise 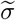 will be unknown in most circumstances, we may treat it as a constant. Consequently, the *detectability* 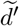 contains all the information necessary to characterize how the detection threshold varies with eye movements.

### III.2. Localization

We also examine another common task of the visual system, *localization*, which tends to accompany detection: if we detect a stimulus, it appears in a specific location in our field of vision. We apply a similar Bayesian framework to quantify how localization is related to detection. Thus, consider a *localization task*, i.e. the task of finding the true location ***l***_0_ of a stimulus within the visual field. While localization accuracy is difficult to measure directly, this task is closely related to a common psychophysical experiment, the *Vernier task*, which requires determination of the *relative* location of two identical stimuli, indirectly quantifying the accuracy of localization. Vernier experiments have shown that the human visual system can resolve offsets smaller than the Nyquist limit set by the cone spacing in the fovea [42–44]. This has been attributed to a variety of different mechanisms [45] including fixational eye movements [46, 47]. In the following, we only consider the continuous localization of a single stimulus. The generalization to a Vernier setup is straightforward but laborious and obfuscates the important observations. A stimulus of known shape, located at the origin, has profile *s*_0_(***r***) with spatial transform *ŝ*_0_(***k***). The same stimulus, relocated at position ***l***, then has the shifted profile

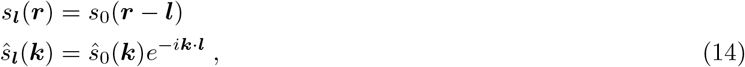

i.e. changing stimulus location gives a phase shift in the signal at each spatial frequency. Thus, instead of deciding between two anticipated expected retinal excitation patterns, localization compares an entire family of patterns, parametrized by the shift ***l***. The IO’s task is to estimate ***l*** on the basis of the received signal *y*(***r***, *t*). In complete analogy to our previous considerations on detection, the posterior probability of a stimulus located at ***l*** is given by

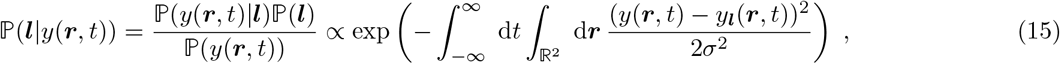

where *y*_***l***_ denotes the excitation pattern generated by a shifted stimulus *s*_***l***_. Here we have assumed “flat” priors such that ℙ (***l***) is independent of ***l*** and the only dependence on ***l*** comes from the likelihood function ℙ (*y*(***r***, *t*)|***l***).

The IO’s estimate for the stimulus location is at the position ***l*** = ***l***_max_ of maximal probability, i.e. the position ***l*** that maximizes the likelihood in eq. (15). Now suppose that the stimulus is actually located at position ***l***_0_. The time-dependent retinal response *y*(***r***, *t*) depends on ***l***_0_ and on the noise *η*(***r***, *t*). Because of the noise, the likelihood will generally be maximized at a position ***l***_max_ unequal to ***l***_0_. There will thus be an error Δ***l*** = ***l***_max_ − ***l***_0_ in the position estimate, but if the signal is strong enough relative to the noise this error will be small. In Appendix B we derive the equation that links ***l***_max_ to any specific realization of noise. In the limit of large signal-to-noise ratio, we then show that the probability distribution for the localization error Δ***l*** takes the shape of a two-dimensional Gaussian distribution with zero mean, i.e. the maximum likelihood estimate ***l***_max_ of location will be distributed symmetrically around the true location ***l***_0_. Figure 2 illustrates the process. The limit of high signal-to-noise ratio ensures that the likelihood distribution does yield a unique maximum. This implicitly confines the range of applicability to |Δ***l***| ≲ *β*. Yet, the analysis of the detection task below shows, if a stimulus can be detected that corresponds to the high signal-to-noise regime in the sense that we use here. The variance of localization error Δ***l*** is characterized by the covariance matrix

**Figure 2.**
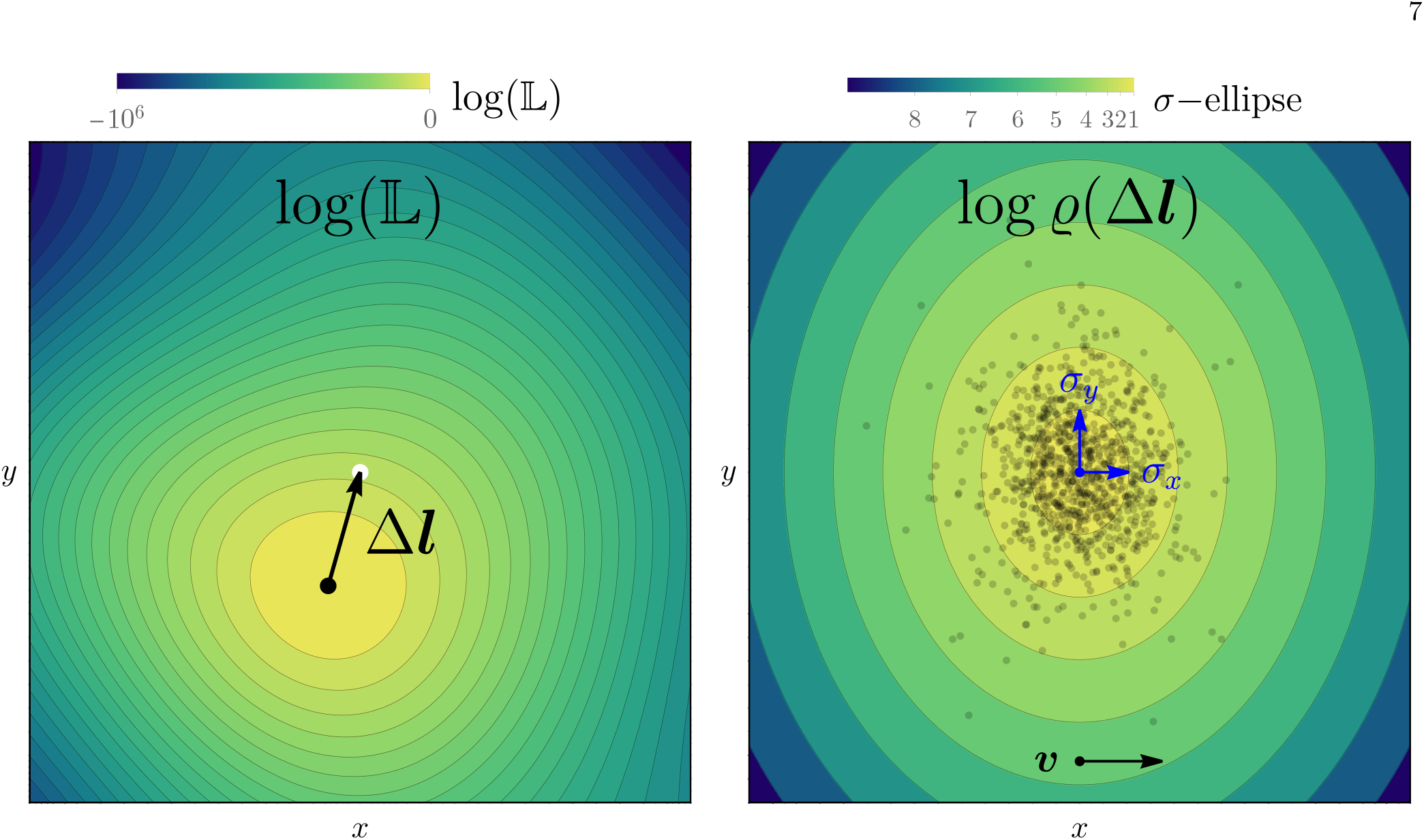
Schematic illustration of localization accuracy of the ideal observer. Left: After sufficient information acquisition the likelihood distribution L of the stimulus location for a specific realization of white noise exhibits a sharp maximum (log-likelihood is shown). The location of that maximum (black dot) corresponds to the maximum-likelihood estimate of the stimulus location but deviates from the actual position of the stimulus (white dot) by Δ***l***. Right: The probability density *ϱ* of the distribution of maximum likelihood estimates of the location approaches a Gaussian as the signal-to-noise ratio increases. Black dots mark simulations of Δ***l*** for different noise realizations and a point like stimulus with straight eye movements in *x*-direction as indicated by the ***v***-arrow. Anisotropic eye movements cause an anisotropy in the probability distribution, i.e. localization accuracy depends generally on direction.

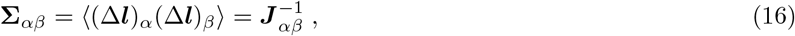

with *α, β* ∈ {*x, y*} which is given by the inverse of the 2 × 2 matrix ***J*** with components

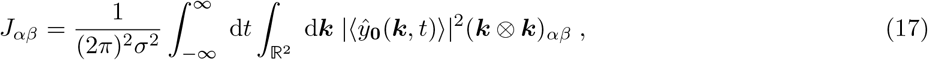

where ⊗ describes the outer product. The localization variance depends exclusively on the noise-averaged excitation pattern generated by the stimulus centered at the origin ⟨*ŷ*_0_(***k***, *t*)⟩ but this function still depends on eye motion.

In contrast to the detection task, a single scalar quantity such as *d*′ is generally not enough to describe the IO’s accuracy of localization. As localization depends on direction, we instead have a two-dimensional tensor. Any anisotropy in the stimulus shape or its blur translates into anisotropy of the accuracy of localization. For example, an oblong rectangular stimulus is typically easier to locate perpendicularly to its long side as more receptors will detect change in response to a corresponding eye movement. Anisotropic eye movements have the same effect, inducing an imbalance between localization parallel and perpendicular to the principal direction of motion. The relevant variance along any specific unit vector ***e*** is given by the scalar product ***e***^*t*^**Σ*e***.

However, if the setup is radially symmetric, i.e. if eye movements are isotropic in an average sense, and the stimulus is also radially symmetric, then *J*_*αβ*_ simplifies to a multiple of the identity matrix ***J*** _*αβ*_ = *l*^′2^1 with a single scalar quantifying accuracy of localization

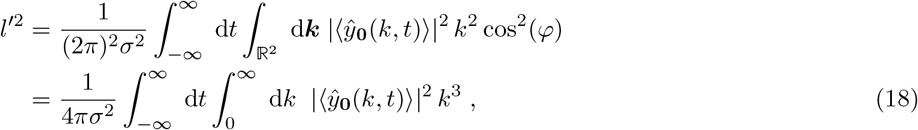

with *k* := |***k***| and polar angle *φ*. Note that the overall variance in the isotropic setup is given by ⟨|Δ*l*|^2^⟩ = 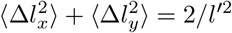, so that large *l*′ corresponds to high localization accuracy.

As we did for the detection task we remove the trivial scaling behavior of *l*′ with noise and contrast, and render the result dimensionless to arrive at the localization equivalent of eq. (10)

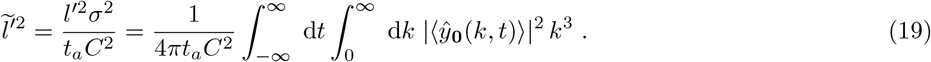

Using the baseline model of section II to express ⟨*ŷ*_**0**_(*k, t*)⟩, this can be cast as

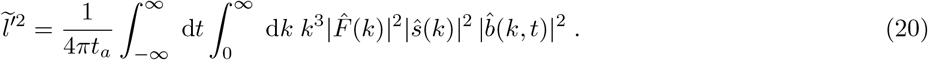

The expressions in eqs. (19) and (20) are strikingly similar to their analogues for detection, i.e. *d*^′2^ of eqs. (10) and (12). Indeed, the only difference, barring inconsequential prefactors, is the factor *k*^2^ arising from the outer product (***k***⊗***k***). Thus, localization along a specified direction becomes theoretically equivalent to detecting the same stimulus with a bias towards higher spatial frequencies. More concretely, the componentwise Fourier transform of the gradient stimulus can be written as ℱ [∇*s*] = ***k****ŝ*(*k*), so that ℱ [∇*s*] ⊗ ℱ [∇*s*] = *ŝ*(*k*)^2^(***k***⊗***k***). That means, detecting the partial derivative of *s*(***r***) in *x*-direction yields an additional factor 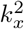 in eq. (11) compared to detecting the stimulus itself. However, the resulting expression is up to prefactors the same as for localizing the original stimulus in *x*-direction. Thus, localization accuracy effectively characterizes the ability of detecting a gradient. This corroborates the intuition that localization is improved by the presence of sharp edges in the stimulus as these lead to larger image gradients which has also been observed experimentally [48]. Conversely, a constant increase in background intensity is easily detected while it is impossible to be localized due to the absence of any spatial variation. This leads to the important theoretical insight: **Localization is equivalent to detection of the stimulus gradient**. This result in itself is independent of fixational eye movements and does not require an isotropic stimulus. Every stimulus has an intrinsic directional “localizability” given by the intensity (squared magnitude) of its corresponding gradient.

A connection between localization accuracy and stimulus gradient has been noted previously in different contexts yet not in relationship with a detection task. In psychophysics it was found that the relevant information for an ideal discriminator in a Vernier task is contained in the first derivative of the stimulus shape [49]. That result is based on the discrimination between two stimuli *s*(*x*) and *s*(*x* + *δx*) and the small *δx* approximation *s*(*x* + *δx*) − *s*(*x*) ≈ *δxs*′(*x*) rather than the posterior probability of the stimulus location for a given noise realization. Another approach utilizes the Cramér–Rao bound that connects the variance of an unbiased estimator to the Fisher information [50]. Indeed, ***J*** as defined in eq. (17) is exactly the Fisher information matrix for the stimulus location. As the distribution of Δ***l*** approaches a Gaussian, the mean offset Δ***l*** becomes an efficient estimator of location for large signal-to-noise ratio yielding the inverse of the Fisher Information Matrix as its covariance. Here, the derivative naturally occurs in the definition of the Fisher information. Since photoreceptors are discrete, the continuous gradient of an image generally does not exist. However, stimuli are typically continuous (on the relevant length scale) and only the stimulus requires differentiation. Thus, even with a discrete receptor array like the human retina, the detectability of the stimulus gradient quantifies localization accuracy.

As the integrand of eq. (20) illustrates, fixational eye movements that control the structure of 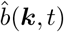 couple to the shape of the stimulus contained in *ŝ*(*k*) in exactly the same way for both tasks, detection and localization. However, since the expressions for detection and localization differ in the factor *k*^2^, the eye can generally not find an optimal eye movement strategy for both detection and localization at the same time. Indeed, below we provide reasons to believe that human eye movements are purposefully optimized for localization accuracy and accordingly sub-optimal for detection.

### III.3. Experimental Protocols: Localization and Detection Thresholds

As with detection, our analysis of localization allows us to determine the accuracy up to constants such as the noise amplitude *σ*, which are unknown or difficult to estimate. However, a measurement of an observer’s ability to localize a stimulus can be referenced to a measurement of their ability to detect it, i.e. their contrast threshold *C*_crit_ (as in eq. (13)). Then, for the localization task, the same stimulus may be presented at a prescribed multiple of the threshold contrast *C* = *αC*_crit_, ensuring that the stimulus is detected consistently. In this case, the accuracy of localization is ⟨|Δ*l*|^2^⟩ = 2/*l*′^2^ with

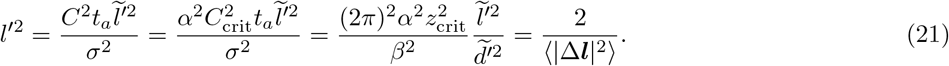

Vitally, the unknown noise amplitude *σ* cancels, so that the final result is independent of this unknown parameter. Thus, under the localization protocol run at fixed multiples of the detection contrast threshold, we are able to predict localization accuracy in terms of the ratio of 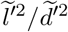, with 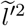 evaluated using eq. (20), and 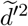 using eq. (11), yielding

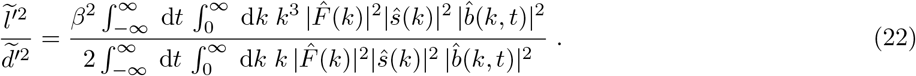

We have assumed radial symmetry throughout for simplicity but eq. (22) is straightforwardly generalized to arbitrary stimuli and eye movements by retaining the full two-dimensional dependence of *ŝ* and 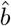 on spatial frequency ***k***. Note that one implicit assumption of the baseline model is that the signal scales linearly with contrast which certainly breaks down for contrasts far above the detection threshold, i.e. *α* ≫ 1. Yet, provided the assumptions of the model hold, the localization and detection tasks do not necessarily have to be performed with the same stimulus presented for the same duration. For example, if the detection task to determine the critical contrast threshold *C*_crit_ is performed for stimulus *s*_*d*_ presented for duration *T*_*d*_, whilst the localization task is performed for *s*_*l*_ presented for duration *T*_*l*_ at a contrast *C* = *αC*_crit_, then the accuracy of localization becomes:

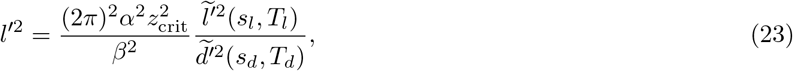

where 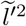 and 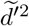 are, as indicated, evaluated for the conditions of the localization and detection tasks respectively.

Localization accuracy and contrast detection thresholds are common experimental observables. However, the ratio 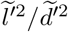 allows us to combine observations from different experiments into a single unified picture of a participant’s visual system. This is a key result, and is one reason why it is important to consider these two tasks within the same framework. In the following we first examine detection and localization tasks in isolation, determining how these tasks are affected by stimulus size, duration, and eye motion. We then proceed to study their ratio 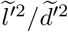.

## IV. MODEL PREDICTIONS

So far we have characterized detection and localization for general linear filters *F* and *K* and arbitrary stimuli and eye movements. In order to provide concrete predictions we need to specify the functional form of *F* and *K*. The characteristic properties of *F* and *K* have been discussed in section II. A detailed description of all the functions chosen for concrete calculations is given in Appendix C. As the stimulus we choose a disk of radius *L*, switched on and off instantaneously according to the amplitude function *A*(*t*) of eq (1).

### IV.1. Diffusive Eye Motion

Fixational eye motions have been characterized using random walks, i.e. a diffusive model [26, 51–53]. Detailed measurements show that eye movements vary substantially across individuals, and a purely diffusive model does generally not provide a fully satisfying description for most people [52, 53]. Nevertheless, in the spirit of utilizing a baseline model with minimal parametrization, the assumption of simple diffusion provides a rough reference for the extent of realistic eye movements, characterized by an effective diffusion constant *D*. For that reason, we compute 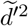 and 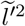 as an ensemble average over all trajectories generated by the same *D*. Calculating the mean square displacement (MSD) of experimental eye traces allows for assigning this diffusion constant to a family of eye movements under the condition that the MSD is found to increase approximately linearly in time. There is some variation in the literature for reported diffusion constants, not least because of variation between individuals, but values of order 10 − 40 arcmin^2^/s are typical [26, 51]. Pure diffusion has the added benefit of being isotropic on average, which coupled to an isotropic stimulus such as a disk of radius *L* allows for a straightforward analysis of 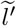 and 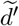 as derived in eqs. (20) and (11). In the following, we map out the impact of diffusive eye motion on detection and localization of a disk. Our goal is to explore how the tasks of detection and localization are affected by random fixational eye movements, characterized by diffusion constant *D*, and by the stimulus, characterized by size *L* and presentation time *T*. In the spirit of our theoretical considerations, we express these parameters in natural units. In the context of visual system response, whether a stimulus is large or small depends on the ratio 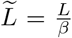. Similarly, whether a stimulus duration is long or short depends on the ratio 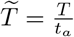. The non-dimensional diffusion constant 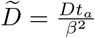 determines whether eye motion is fast or slow in the context of other elements of the visual system: a value 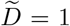 corresponds to the eye moving roughly one blur length in one adaptation time.

With that, the dimensionless measures of detection capability, 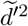, and localization accuracy, 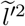 are functions only of the parameter triplet

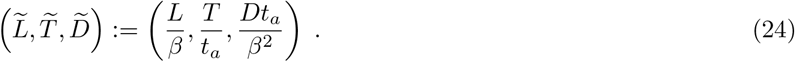

By varying these three quantities, we find regions where 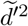 and 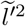 behave in different ways. Fig. 3 illustrates the typical transitions for detection of a small stimulus, i.e. 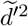 for 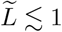. The behavior is different depending on whether the scaled diffusion constant is large or small.

**Figure 3.**
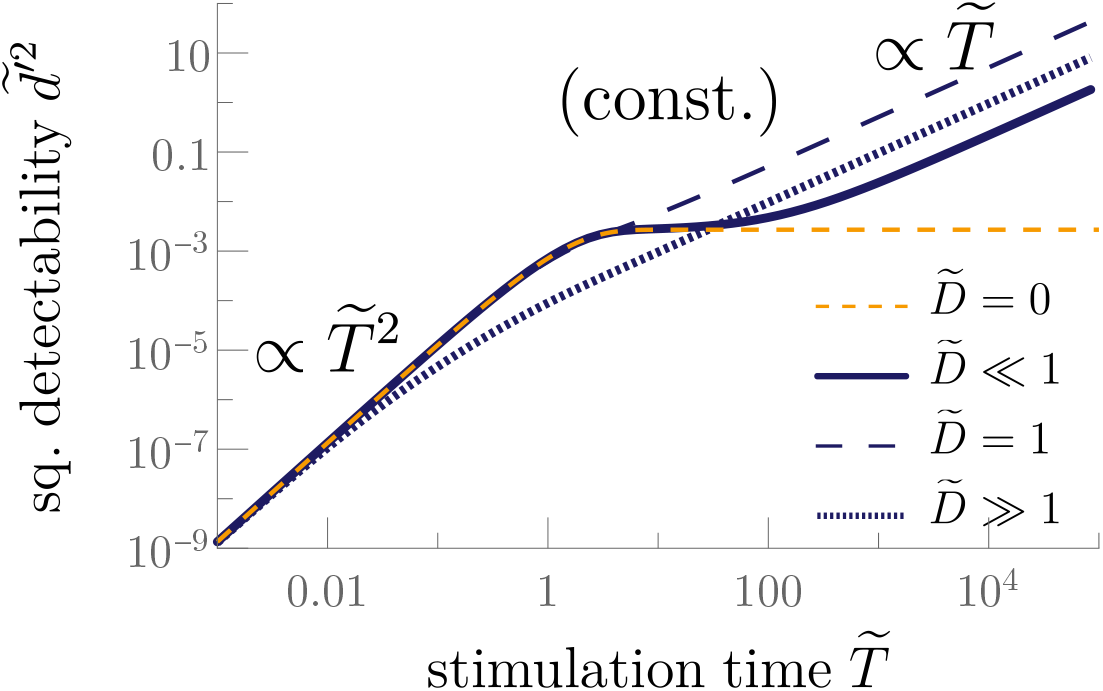
Scaling behavior of the squared detectability 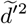 of a small stimulus 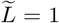 as a function of stimulus duration 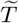 for different diffusion constants 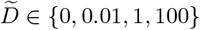.

For small 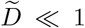 there are three regimes. For brief stimulations 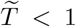, the stimulus is presented faster than the timescales of the temporal filter. As a result, the amount of signal integrated by the temporal filter is proportional to 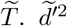 depends on the square of the temporal filter output and thus scales quadratically with presentation time, i.e. 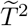. This is Bloch’s law [29, 54] and it occurs for all 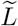 and 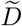 at short presentation times. For 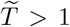, 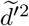 reaches a plateau because the retina adapts to the presence of the stimulus, which has still not moved substantially from its original position. However, when 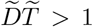 (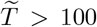 for the solid curve in Fig. 3) the eye motion during the presentation time moves the stimulus a distance further than the blur length, so that the stimulus is presented to new regions of the retina which have not fully adapted to it. As a result, the average signal becomes a steady-state transient and thus independent of time. Integrating this constant excitation pattern over *T* yields a linear increase of 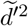 with time. Without eye movements, i.e. 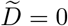, the transient vanishes due to perfect adaptation.

For large 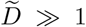, the eye motion is sufficiently fast that the long time transient is reached before adaptation can occur. Hence, there is no adaptation plateau, and 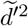 transitions directly from quadratic to linear behavior near 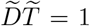 (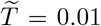 for the dotted line in Fig. 3). However, rapid motion leads to a loss of information for shorter stimulus durations as individual photoreceptors are not exposed to the stimulus for the full excitatory period of the adaptation kernel. At 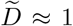 these losses are minimized while maintaining enough motion to avoid the inhibitory part of the kernel resulting in optimal detectability for short and long stimulation times.

The above example illustrates that, for a small stimulus 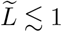, it is possible to identify regions in the two-dimensional space 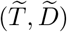 for which fundamentally different behavior of 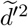 can be observed and we could draw a similar diagram for the squared localizability 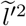. The transitions between different scaling regimes can be associated with a change in the order of the fundamental length scales:

i. the blur length *β*
ii. the average distance traveled during adaptation time 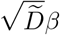,
iii. the average distance traveled during stimulation 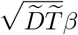, and
iv. (iv) the stimulus size 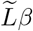.

If the stimulus is small as in Fig. 3, the effective stimulus size is set by the blur so that the actual stimulus size does not play a role. The three-dimensional space of 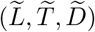 conveniently separates into two regions, small stimuli 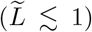 where the stimulus size is smaller than the blur scale and thus negligible, and large stimuli 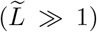. Note that, because of our use of natural units, the terms “small” and “large” are always relative to the blur length *β*, i.e. a small stimulus becomes large just by reducing the blur as shown in Fig. 4. Hence, small stimuli are more intuitively labelled blurry in contrast to large stimuli which are crisp. The size of the stimulus crucially also affects the relationship between stimulus and gradient. The directional derivative of a large stimulus is limited to its perimeter while the gradient of a blur-dominated stimulus covers the same area as the stimulus itself (cf. Fig. 4).

**Figure 4.**
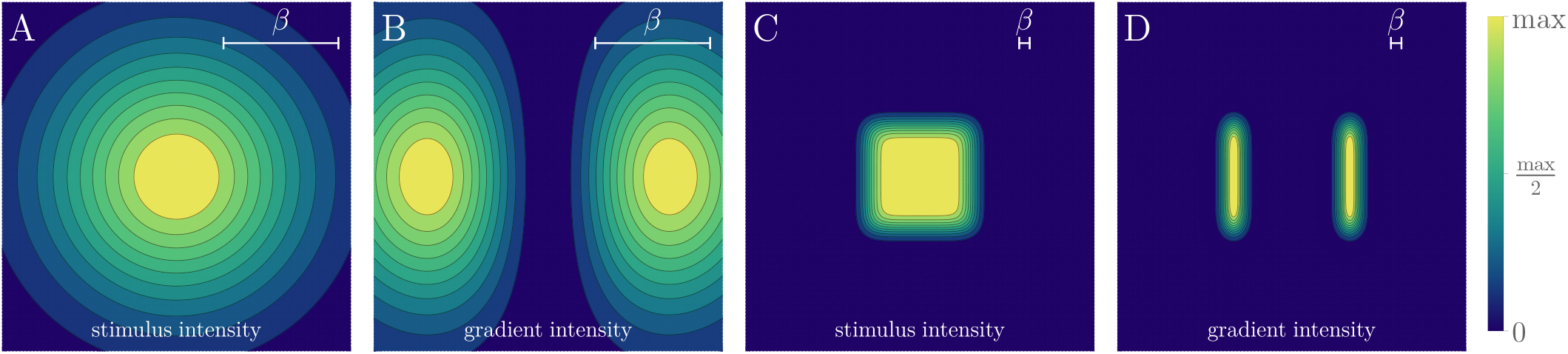
The difference between a “small” stimulus *L* ≲ *β* and a “large” stimulus *L* ≫ *β*: intensity of a unit square stimulus with blur *β* = *L* (A) and intensity of its derivative in horizontal direction |∂_*x*_*s*(***r***)|^2^ (B). This is juxtaposed with the intensities for the same stimulus with less blur *β* = 0.1*L* in (C) and (D). Each intensity is normalized to its maximum to emphasize its spatial extent. The term small refers to the rescaled length 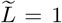 yielding a blurry stimulus (A, B), while a large stimulus 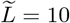 is crisp (C, D). The intensity of a large stimulus scales with its area (C), while the intensity of its gradient scales with the stimulus perimeter (D). In contrast, for a stimulus dominated by blur, stimulus and gradient intensity scale in the same way with stimulus size, that is with 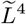 (A, B).

Fixing the stimulus size as small or large allows us to produce 2D diagrams in 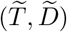 for either stimulus size separately, covering the complete phenomenology of the three-dimensional parameter space. Figures 5 and 6 present the corresponding maps of this 2D space, outlining the regions with different asymptotic scaling regimes separated by white lines, together with a contour plot of 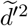 and 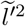. We refer to such a plot as a scaling diagram. Scaling diagrams allow us to communicate compactly all the different scaling regimes and their corresponding transitions for both detection 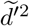 and localization 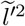.

**Figure 5.**
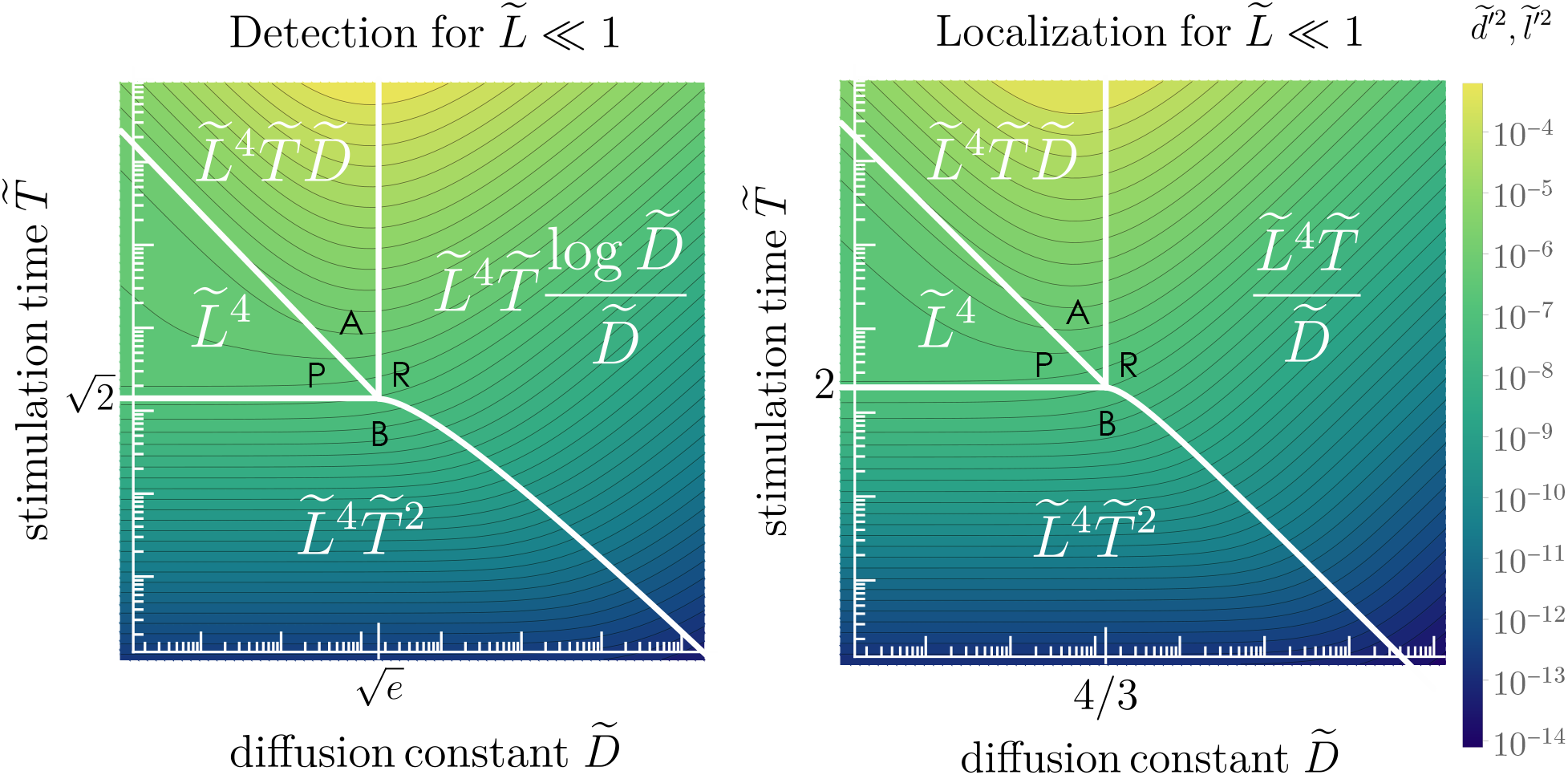
Logarithmic contour plots of the squared detectability 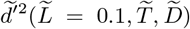 (left) and the squared localizability 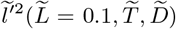 (right) for a small stimulus highlighting the transition between different scaling regions. The leading order scaling is indicated in white, the black letters are labels to simplify the reference to the text. The coordinates of the “quadruple point” where all scaling regimes meet is independent of 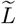 to first order, its coordinates for a point-like stimulus are marked on the axes. Large axis ticks mark powers of 10. Vertical slices through the left diagram generate the curves shown in Fig. 3. The legend applies to both plots and contours are equally spaced on the logarithmic scale.

**Figure 6.**
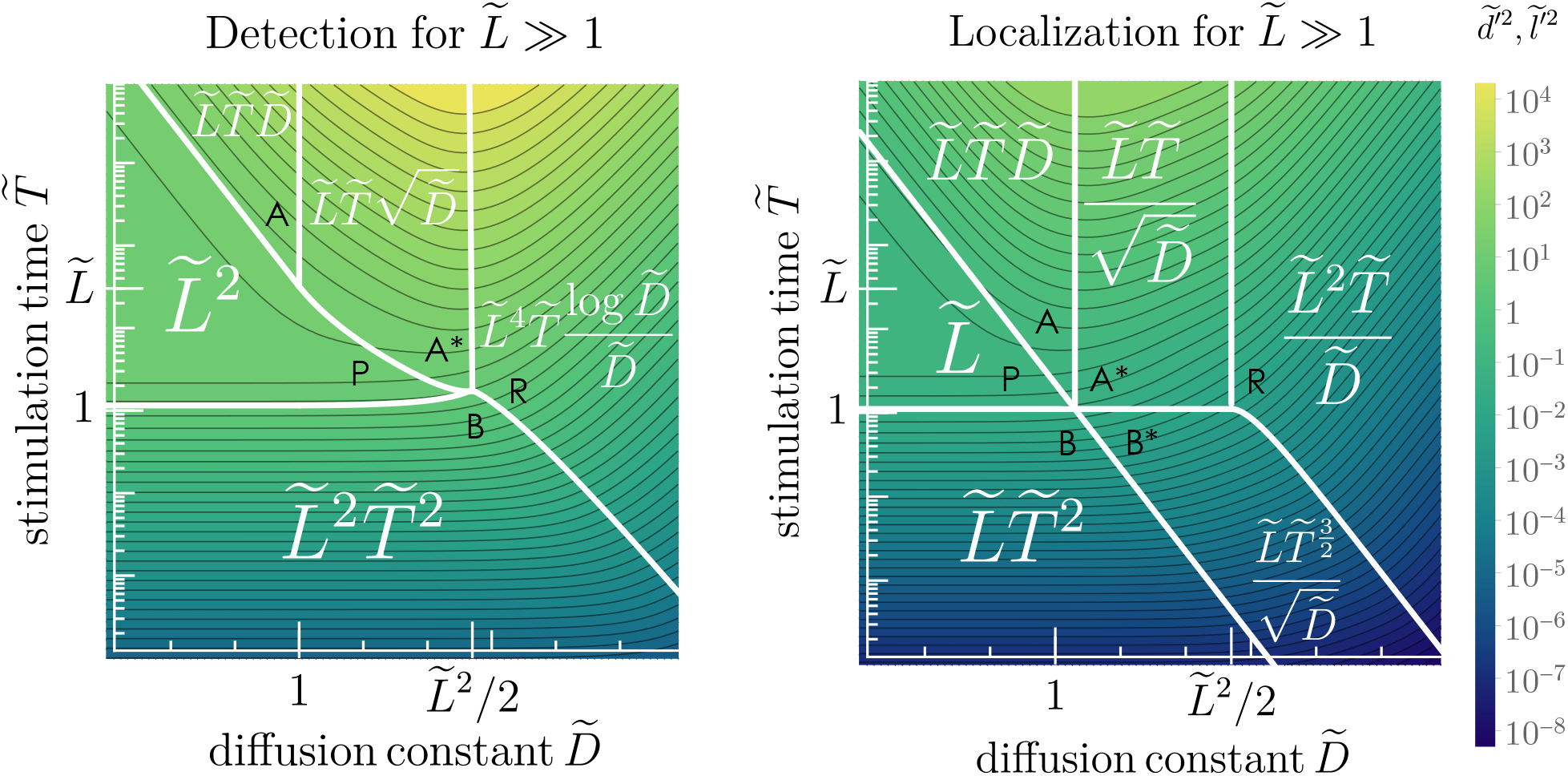
Logarithmic contour plots of the squared detectability 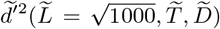 (left) and the squared localizability 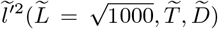 (right) for a large stimulus highlighting the transition between different scaling regions. The legend applies to both plots and contours are equally spaced on the logarithmic scale.

### IV.2. Small Stimuli

For 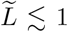, the stimulus is smaller than or comparable to the blur scale, so that the retinal image is dominated by blur. If the primary source of blur is set by the cone spacing and the eye’s optics, then an object that generates a retinal image of this angular size is either extremely small or exceptionally far away. For comparison, stars in the night sky appear to cover roughly 60” due to atmospheric blur, giving 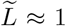. Hence the condition 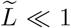 is rarely encountered in natural viewing.

Fig. 5 presents scaling diagrams for both detection 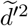 and localization 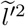 for a stimulus small compared to the blur size. The scaling with 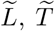 and 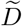 is indicated in each region, delineated by white boundaries. In this small stimulus limit, the scaling with 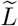 is consistent across the whole diagram. The signal integrated by the spatial filter is proportional to the stimulus area, 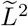, and since both 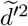 and 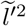 depend on the square of the output signal, this results in an overall scaling of 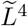. To structure the discussion of the scaling with 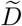 and 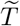 we refer to the black letters in Fig. 5.

#### Detection

Detection is characterized by 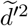, shown in the left panel of Fig. 5:

(B – Bloch Regime): For 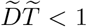, the eye motion is sufficiently slow, relative to the stimulus presentation, that the effects of motion are negligible, and hence the result is independent of 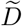. This gives a scaling of 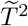 (Bloch’s law) for 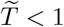 as described above for Fig. 3.

(P – Plateau Regime): For 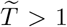 but 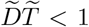, the eye does not move enough within one adaptation cycle for the stimulus to exit the patch of retina that it initially fell on. Consequently, signal is lost due to adaptation and a plateau emerges.

(A – Asymptotic Transient Regime): For 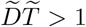 and 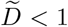, eye motion during stimulus presentation has a significant effect, because the stimulus is presented to multiple blur-sized patches of the retina, thus partially overcoming adaptation. A steady transience emerges giving rise to a linear dependence of 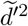 on presentation time 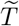. This transience is maximized for 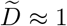. Increasing 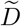 up to 1 increases the signal that passes through the temporal filter so that the detection capability 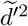 increases with 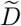.

(R – Rapid Movement Regime): If motion is quicker than adaptation, i.e. 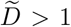 and 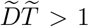, the time spent on each patch of retina becomes shorter than the adaptation time, so that the accumulated signal on each patch is sub-optimal. Hence, 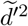 decreases with 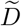 for any given 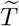 the more rapid the eye motion becomes.

One important feature of performance that emerges from this analysis is that for short presentation times relative to adaptation, eye motion is either inconsequential or detrimental.

The optimal motion, giving rise to maximal 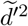 at each presentation time for 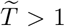, and negligible detriment for 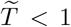, is at 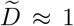, so that the eye moves approximately one blur length per adaptation time. For a point-like stimulus in the limit 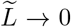, all scaling regimes meet at the “quadruple point” 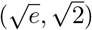. Correction terms arise for a finite-size stimulus which slightly shift the scaling transitions which are broad and continuous anyway. Thus, we neglect stimulus-specific coefficients in the discussion, treating all numerical constants as 1.

#### Localization

The right panel of Fig. 5 shows the scaling diagram for localization, 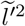. This underlines that, for small stimuli, detection and localization are almost indistinguishable. This is expected because of the similarity in scale of the stimulus and its gradient (Fig. 4 A and B). Thus, all explanations for the detection diagram carry over to localization. The regime transitions shift slightly but the structure of the scaling diagram remains the same. The “quadruple point” for a point-like stimulus is found at (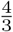, 2) which again can comfortably be treated as of the order 1. The only scaling difference arises for large diffusion constants. The dilution of the excitation pattern due to very extensive eye movements is slightly less harmful for detection compared to localization due to an additional logarithmic dependence on 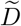.

In summary, **for a small stimulus, detection and localization vary identically with stimulus size, duration, and eye motion**. With detection and localization effectively being the same task, the two operations can be optimized simultaneously. This has practical relevance outside human vision, as for example in single particle tracking. Phase masks optimized for localization of fluorescent emitters which are effectively point-like stimuli also optimize detection probability [55]. Other examples can be found in astronomy, where matched-filter detection is optimal for single-image point source detection while also saturating the Cramér-Rao bound [56].

### IV.3. Large Stimuli

Most real-life stimuli are larger than the blur length. This introduces 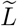 as additional relevant length scale and leads to richer scaling diagrams, presented in Fig. 6.

#### Detection

For detection 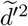 (Fig. 6, left) the general form of the temporal scaling remains the same: a quadratic regime for short times, a plateau if diffusion is slower than adaptation, and a linear regime corresponding to the long time transient. However, the scaling of each regime with stimulus size is substantially different from the small-stimulus case.

We consider initially the case that the eye motion is not so rapid as to move the stimulus a distance comparable to its own size during the presentation, i.e. 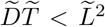 (P and B). For short presentation times 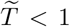 (B) the stimulus is flashed over an area of retina proportional to 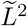, so that Bloch’s law gives 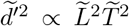. For longer presentation times, 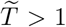, the retina adapts during the stimulus presentation, so that in the absence of eye motion 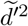 plateaus to a value proportional to the area of the stimulus on the retina 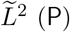.

As before, we expect eye motion to overcome adaptation for 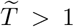. However, if 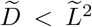 (A and A^∗^), the eye moves less than the stimulus size during one adaptation time, so that the retinal signal induced by eye motion is confined to a band around the perimeter of the stimulus. The width of this band is either the blur scale *β* (for 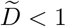) (A) or the distance 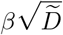 moved during one adaptation time for 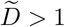 (A^∗^). This has two consequences:

Firstly, the contributions to 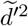 from eye motion are proportional to the stimulus perimeter, i.e. 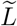. However, contributions to 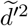 from stimulus modulation (switching the stimulus on and off) are proportional to stimulus area, i.e. 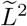. Thus, the detectability is dominated by the initial turning on of the stimulus, and it is harder for eye motion to provide significant additional evidence. As a result, it takes longer presentation times before the effects of eye motion are observed, and the range of presentation times for which there is an adaptation plateau increases with 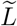 for any given diffusion constant. This can be seen in Fig. 6: For 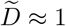 the adaptation plateau occurs for presentation times 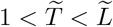. Decreasing 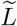 reduces the duration and magnitude of the adaptation plateau, until for small stimuli (Fig. 5) the adaptation plateau vanishes entirely at 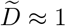. This is exactly the behavior observed by Barlow [27, 57], who measured contrast thresholds for large and small stimuli, observing an adaptation plateau for a large stimulus but not a smaller stimulus.

Secondly, for both regimes, (A) and (A^∗^), the retinal signal due to eye motion, summed around the perimeter of the stimulus, increases with increasing diffusion constant, albeit more slowly in (A^∗^). Consequently, the optimal motion for stimulus detection is at 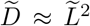, maximizing 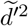 at each presentation time for 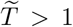 at a negligible detriment for 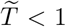. It is also the smallest diffusion constant for which the adaptation plateau disappears. This is due to the eye moving a distance of order the stimulus size in one adaptation time, so that the stimulus falls on an entirely new patch of retina during each cycle of adaptation. That means, **for a detection task the optimal eye motion depends on the scale of the stimulus**.

For 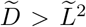 and 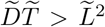 (R) the eye moves further than the stimulus size within both adaptation and stimulus presentation time. As with the small stimulus, the accumulated signal on each patch of retina becomes sub-optimal, and 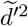 decreases as 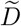 increases further. The stimulus is dynamically blurred by the movement and thus is effectively a small stimulus relative to blur, i.e. the scaling coincides with the R-regime of a small stimulus.

#### Localization

The localization scaling diagram for large stimuli is presented in the right panel of Fig. 6 and is substantially different to the corresponding diagram for detection. This is primarily due to the scaling with stimulus size. As established above, localization is equivalent to the detection of the stimulus gradient (cf. Fig. 4). Thus, only the edges of the stimulus contribute to localization. For a disk stimulus that means, as long as diffusion during both adaptation time and stimulus duration is small compared to the stimulus size, only the perimeter of the stimulus provides information and thus 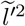 scales only linearly with 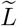 for the majority of the scaling diagram (all except (R)). In contrast to the detection task, this also applies for short presentation times and the adaptation plateau, because the stimulus gradient vanishes in the center of the disk even without adaptation.

Since the contributions to 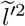 from stimulus modulation are proportional to stimulus size 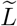, size provides relatively less information in support of stimulus localization as compared to detection. For localization, the adaptation plateau disappears for 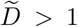 similar to the small stimulus case and the plateau value is reduced compared to detection. Due to the reduced plateau level, information gained by eye motion during extended stimulation is more significant for localization compared to detection.

Accordingly, the optimal diffusion constant for stimulus localization is at 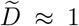, giving rise to maximal 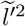 at each presentation time for 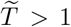 with negligible detriment for 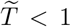. Again, optimal movement corresponds to the slowest motion for which the adaptation plateau disappears. This is because the sharp stimulus gradient at the perimeter of the disk is convolved with the blur length of the spatial filter. As for the small stimulus case, the optimal motion corresponds to the eye moving a distance of order the blur length in one adaptation time. For small diffusion constants (B, P, A), localization of a large stimulus scales with 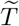 and 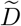 in exactly the same way as for localization of a small stimulus.

For larger diffusion constants, the localization scaling diagram for large stimuli differs in detail from the small stimulus case, as the eye motion interacts with the scale of the whole stimulus. Notably, for 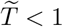 and 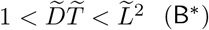 (B) an additional 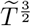 scaling regime appears. Here, diffusion distance during the stimulus presentation time is larger than the blur scale but smaller than the stimulus scale. This additionally blurs the stimulus gradient so its effective width is given by 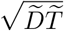. This dynamic blur reduces the intensity of the excitation pattern by a factor 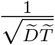 compared to Bloch’s regime giving rise to the peculiar 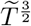 scaling. More generally, for 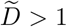 and 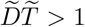 (A ∗, B ∗, R) the eye moves further than the blur length within both adaptation and stimulus presentation times, whilst for 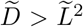 and 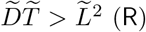 it also moves further than the stimulus size. The accumulated signal on each patch of retina becomes increasingly sub-optimal, and 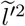 decreases as 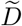 gets bigger.

A key conclusion from the analysis of the scaling diagrams is that, for large stimuli, the optimal diffusion constants for localization and detection are separated by a factor of 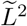 which can account for several orders of magnitude. Thus, a diffusive eye cannot simultaneously optimize motion for detection and localization of large stimuli. Correspondingly, their scaling diagrams do not match giving rise to regimes where localization is significantly improving with increased presentation time whilst detection is not. This is of immediate experimental interest.

### IV.4. Localization at Detection Threshold

As noted in section III.3, we now utilize our theoretical framework to characterize localization accuracy at the detection threshold, the key quantity being the dimensionless ratio 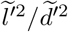. This quantity has the advantage that it is independent of the unknown noise parameter 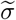 allowing for absolute predictions. As it establishes a fundamental link between two common psychophysical observables, i.e. detection thresholds and Vernier acuity this quantity bears additional practical relevance. Within our model assumptions the potentially non-monotonic way that 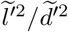 changes with stimulation time for a fixed stimulus size depends exclusively on the interplay between FEMs and the spatial and temporal filters. That provides a straightforward experimental protocol to study the behavioral implications of fixational eye movements. Having examined localization and detection separately, we simply need to take the ratio of the corresponding scaling diagrams. Fig. 7 illustrates the results for both small and large stimuli.

**Figure 7.**
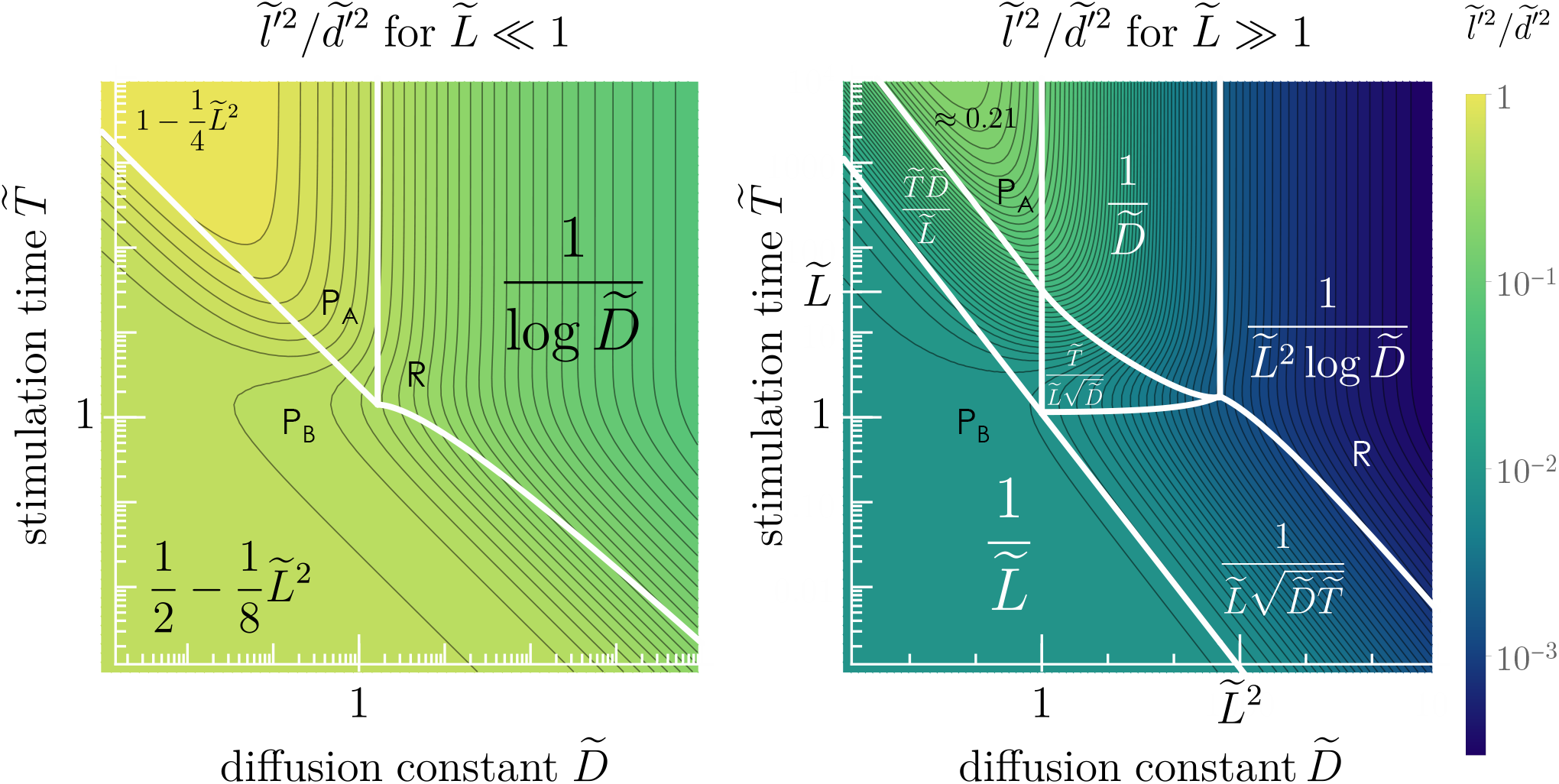
Localization accuracy at detection threshold: Contour plots of the ratio 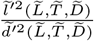 for a small stimulus 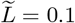 (left) and a large stimulus 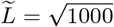 (right). Both diagrams use the same logarithmic color function and contour lines. For small stimuli, the ratio does not exhibit scalable effects. For large stimuli, a plethora of different scalings emerges which can be utilized experimentally.

For small stimuli, the ratio of localization and detection is relatively flat, emphasizing the effective equivalence of both tasks for blur-dominated stimuli. Although 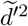 and 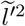 each individually cover 10 orders of magnitude in the chosen parameter range, their ratio varies just above 1 order of magnitude. The experimentally accessible parameter range is even more confined. For small eye motions we find a plateau at short times with the ratio 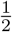 for the point-like stimulus (P_B_) because 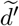 and 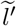 scale identically in the Bloch regime. For longer times, a second plateau emerges separated by a factor of 2 that corresponds to identical scaling in the asymptotic transient regime (P_A_). Absolute accuracy for localization at the detection threshold is maximal for a point-like stimulus observed for long times at 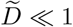. The limiting factor is the blur length capping the resolution at 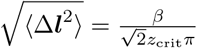 which for a reasonable blur length of 1′ at a threshold *z*_crit_ = 3 corresponds to roughly 4.5″. However, a flashed point can already be localized with a standard deviation 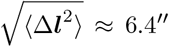. Thus, a small stimulus becomes only slightly easier to localize at the detection threshold if it is presented for a longer time and significantly smaller diffusion constant than optimal for either localization or detection. Eye movements too extensive likewise hurt performance in either task but localization suffers slightly more in comparison (R).

For large stimuli (right panel of Fig. 7) the scaling diagrams for detection and localization differ substantially, so there is significant variation in localization at detection threshold across the explored parameter space. Most prominently, for short presentation times, detection 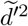 scales as stimulus area, i.e. 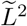, whilst localization scales with stimulus perimeter 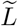. Accordingly, the Bloch plateau (P_B_) decreases with stimulus size. This regime is easily accessible experimentally and the effect depends exclusively on the properties of the stimulus. As this arises entirely due to the spectral relationship between the stimulus and its gradient, spatial and temporal profile of the stimulus could be chosen to emphasize the effect. A brief flash of a large stimulus is easily detected but impossible to locate accurately.

Further, there is significant effect of stimulus presentation time. Close to the optimal diffusion constant for localization 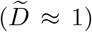, the ratio 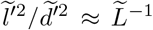 is constant for short presentation times, but increases beyond 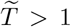 until reaching another constant value 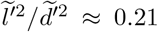 for 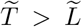 which is independent of the stimulus size (P_A_). Thus, the localization accuracy at the detection threshold varies significantly with stimulus presentation time, even non-monotonically for mesoscale eye movements 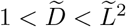.

Remarkably, eye movements faster than the blur length 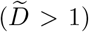 are universally detrimental for localization while detection still benefits from more exploratory eye movements. As a consequence, an ideal observer with excessive eye movements should have comparatively larger Vernier thresholds but smaller detection thresholds if the respective stimuli are large compared to the blur scale.

### IV.5. Explicit Predictions

So far, all our results were presented in natural units. To get a feel for the actual magnitude of the effects of eye movements we present explicit predictions for a visual system characterized by reasonable parameters such as *t*_*a*_ ≈ 40 ms and *β* ≈ 1 arcmin. Diffusion constants have been reported for FEMs in the range 0.01 − 0.04 arcmin^2^*/*ms [26, 51]. Notably, the natural scale of the diffusion constant corresponding to 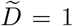 is given by *β*^2^*/t*_*a*_ ≈ 0.025 arcmin^2^*/*ms so that real eye motion seems to roughly have the extent of one blur length per adaptation time. Thus, natural eye motion appears to be tuned towards optimality for localization (which corresponds to 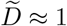) of any stimulus, rather than for detection of large stimuli. For realistic parameters we now consider a set of experiments aimed towards resolving the impact of FEMs on test performance.

#### Temporal and Spatial Summation

The absolute detection threshold depends on the unknown receptor noise parameter 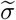. However, the critical contrast is proportional to 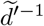. That means, irrespective of the absolute contrast corresponding to the threshold, we expect qualitatively different behavior of the threshold for small and large stimuli as demonstrated in Fig. 8 (left). For short stimulus presentations, the Bloch regime causes detection thresholds to improve linearly with time irrespective of the stimulus size. However, after one adaptation cycle, a large stimulus falls primarily on patches of the retina that have adapted to the stimulation. As a consequence, the contrast threshold plateaus for as long as the eye movements need to transfer the stimulus to a fresh part of retina and the duration of the plateau grows linearly with the stimulus size *L*. For a small stimulus, the plateau vanishes as the detectability transitions directly from the Bloch regime to the asymptotic asymptotic transient regime. Temporal summation in this regime depends on the type of stimulus. For the disk stimulus, the contrast threshold decreases with the square root of the stimulus duration.

**Figure 8.**
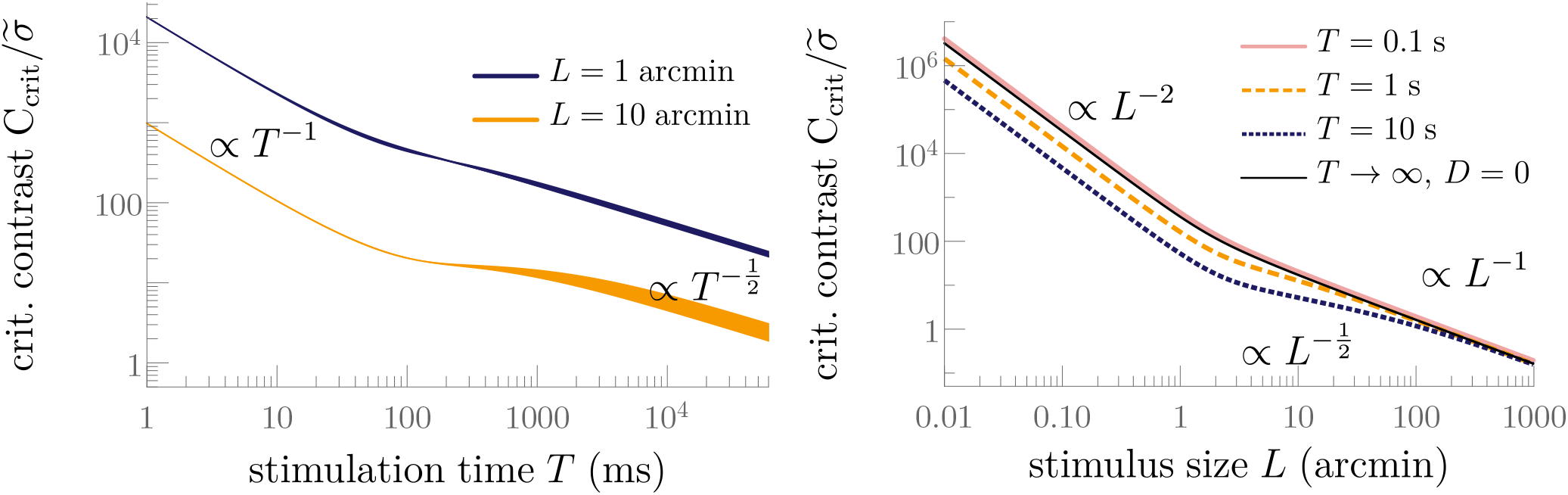
Left: Different behavior of the detection threshold for small as opposed to large stimuli. The thickness of the plots corresponds to the range of diffusion constants 0.01 – 0.04 arcmin^2^*/*ms. The large stimulus always has the lower detection threshold but it reaches a plateau level after 100 ms whereas the threshold of a small stimulus keeps decreasing with the square root of stimulation time. The plateau lasts for the time it takes the eye to diffuse a distance of the order of the stimulus size. Right: At the blur length, the detection threshold changes its scaling behavior with *L*, transitioning from the small stimulus to the large stimulus regime. This effect does not depend on FEMs, but also exists without eye movements *D* = 0. The colored curves use *D* = 0.025 arcmin^2^*/*ms.

With fixations of a naive observer rarely lasting longer than 500 ms, the time window available to distinguish time dependencies is limited. Thus, while the plateau can be extended by increasing the stimulus size, only the beginning of the plateau is easily accessible experimentally. Beyond 30 arcmin the stimulus becomes larger than the foveola so that center and perimeter of the stimulus may fall on patches of retina with vastly different densities of cone photoreceptors [36]. Under these circumstances our retinal continuum simplification becomes increasingly questionable. Importantly, for stimulus presentation *T* ≲ 100 ms, the contrast threshold is largely independent of movements. Thus, only between 100 ms and 500 ms there is a scaling difference to resolve. For a small stimulus, the detection threshold should improve by a factor of 2 in this time frame. However, due to the transition to the plateau even for the large stimulus, the threshold is expected to improve by 25%. Thus, the effect is comparably small and it is advised to use stimulus modulation such as a slow turning on and off of the stimulus to reduce the impact of the initial transient. Without such measures, the majority of information is integrated before adaptation becomes relevant so that the trajectory of the eye beyond 2*t*_*a*_ hardly affects the contrast threshold.

Nevertheless, the difference in scaling behavior between small and large stimuli has been observed experimentally. Barlow measured increment detection thresholds in the periphery at 6.5^°^ eccentricity [57]. The detection of a large stimulus decreases linearly during the first 100 ms indicating complete summation before transitioning to a plateau corresponding to zero summation. In contrast to that, the detection threshold of a small stimulus continues to decrease for stimulations longer than 100 ms but at a reduced rate consistent with a square root dependence as pointed out by previous work [27]. However, as the accessible data hardly spans a decade in time and threshold contrast, the square root dependence is intrinsically hard to resolve. The gradual transition from complete to partial summation further complicates the extraction of power law exponents.

Perhaps more straightforward is to vary the stimulus size at a constant stimulation time as exemplified in the right panel of Fig. 8 to study spatial summation. The critical contrast exhibits a scaling transition around the blur length which could be interpreted as the size of the equivalent receptive field that combines signals spatially early in visual processing.

For small stimuli 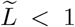, the detectability 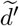 scales with 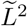 irrespective of eye movements or stimulation time. In the classical literature, it is more commonly expressed in terms of the stimulus area *A*_*s*_ and known as Ricco’s law of complete spatial summation, i.e. 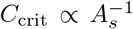 [57, 58]. The area corresponding to the upper limit of complete summation is commonly referred to as Ricco’s area and its dependence on stimulus duration, background intensity, and retinal stabilization has been the subject of numerous studies (e.g., [57–62]).

For large stimuli, the scaling with 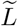 depends on the scaling regime. In the Bloch regime and in the plateau regime the detectability 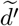 scales linearly with 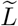 which translates to Piper’s law, i.e. 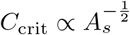 [63]. Irrespective of the stimulus duration, there is always a transition around the blur length corresponding to the change from a small to a large stimulus. More precisely, without eye movements (black line in Fig. 8) the asymptotes can be determined analytically and they intersect at *L*_ricco_ = 2*β* (radius). Thus, blur lengths associated with pre-neural mechanisms (0.5’ - 1.0’) result in Ricco’s areas between *π* arcmin^2^ and 4*π* arcmin^2^. Our model ultimately does not distinguish whether this area describes the pre-neural [60] properties of the fovea or is the result of post-receptor pooling [62]. However, associating *β* with the pre-neural length scales such as cone diameter leads to predictions of Ricco’s area consistent with the experimental measurements in the fovea at low photopic conditions [62, 64].

For stimulation times 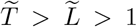, we enter the asymptotic transient regime which for large stimuli scales only with the square root of the stimulus size (cf. Fig. 6 showing 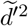). Thus emerges an intermediate range of stimulus sizes for which the detection threshold scales like 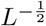. This corresponds to a fourth-root law in the stimulus area 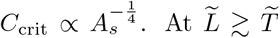, we fall back into the plateau regime and hence again find Piper’s law for large stimuli. The fourth-root law is strongly constrained by the stimulus duration – resolving it cleanly requires stimulations of several seconds. Yet, even after one second of stimulation, the deviation from a model without eye movements (black line in 8) is noticeable. That bears particular relevance because the regime beyond Ricco’s law ceases to be a single power law. Consequently, power-law fits aimed at extracting a single exponent will be systematically off and could be mistaken for Piper’s law with exponents larger than the expected 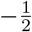 (in the stimulus area). Fourth-root spatial summations have been hypothesized previously based on experimental observations [65, 66] and explained by a “noisy energizer” model [67] which parallels the derivation of Piper’s law but with a squared contrast transducer. Importantly, all the studies finding approximate fourth-root dependencies were conducted with stimuli consisting of various types of gratings, i.e. stimuli that feature spatial variation over small distances. That means fixational eye movements of half a cycle per adaptation time are sufficient to avoid the plateau regime, extending the range of the asymptotic transient regime substantially. This would explain why the fourth-root law has only been reported for grating stimuli and suggests fixational eye movements as the key mechanism. In any case, our analysis illustrates that fourth-root spatial summation does not require an energy (squared contrast) transducer. The existence of the fourth-root regime underlines one strength of our analytic approach, as it allows us to study stimulus parameters that highlight the existence of a performance regime which would be easy to overlook in numerical simulations.

#### Localization at Detection Threshold

Finally, we combine our findings on detection and localization, asking how well a stimulus can be localized if an observer can just barely detect it. Eq. (21) directly links the ratio 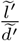 depicted in Fig. 7 to the localization accuracy at the detection threshold. As the detection threshold varies with stimulus size and duration, the corresponding localization task needs to be performed with a variable contrast adjusted to the detectability of the stimulus. Experimentally, that requires the determination of the detection threshold separately for each stimulus condition followed by a localization task at the corresponding contrast (or fixed multiples thereof). By choosing the contrast for the localization task relative to the detection capability of the observer, the localization accuracy does not depend on the unknown internal noise parameter *σ* any more. Within our model, the localization accuracy at the detection threshold depends exclusively on the observer’s fixational eye movements. In Fig. 9 (left) we show predictions of localization accuracy at detection threshold for diffusion constants associated with realistic FEMs 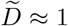. There are two key differences between localization and detection for large stimuli: First, the scaling with stimulus size in the Bloch regime and second, the existence of a plateau regime for detection. Thus, localization accuracy at the detection threshold is constant for the duration of the Bloch regime at a value that grows proportional to 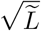. For observation times beyond the Bloch regime (*T* ≳ 80 ms), localization at threshold improves with longer stimulus presentations. However, this decay is confined to a time window of duration 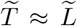. With the regime so narrowly confined the theoretical 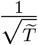-behavior can only be observed for very large stimuli. For a 10’ stimulus, the decay lasts merely about 400 ms and is thus dominated by the transitions to the short- and long-time asymptotic plateau values, respectively. The long-time asymptotic is technically independent of the stimulus size but it varies roughly by a factor of 2 between the class of small stimuli 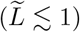 and large stimuli 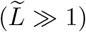 as indicated by the slightly different constants in Fig. 7 in the asymptotic transient regime.

**Figure 9.**
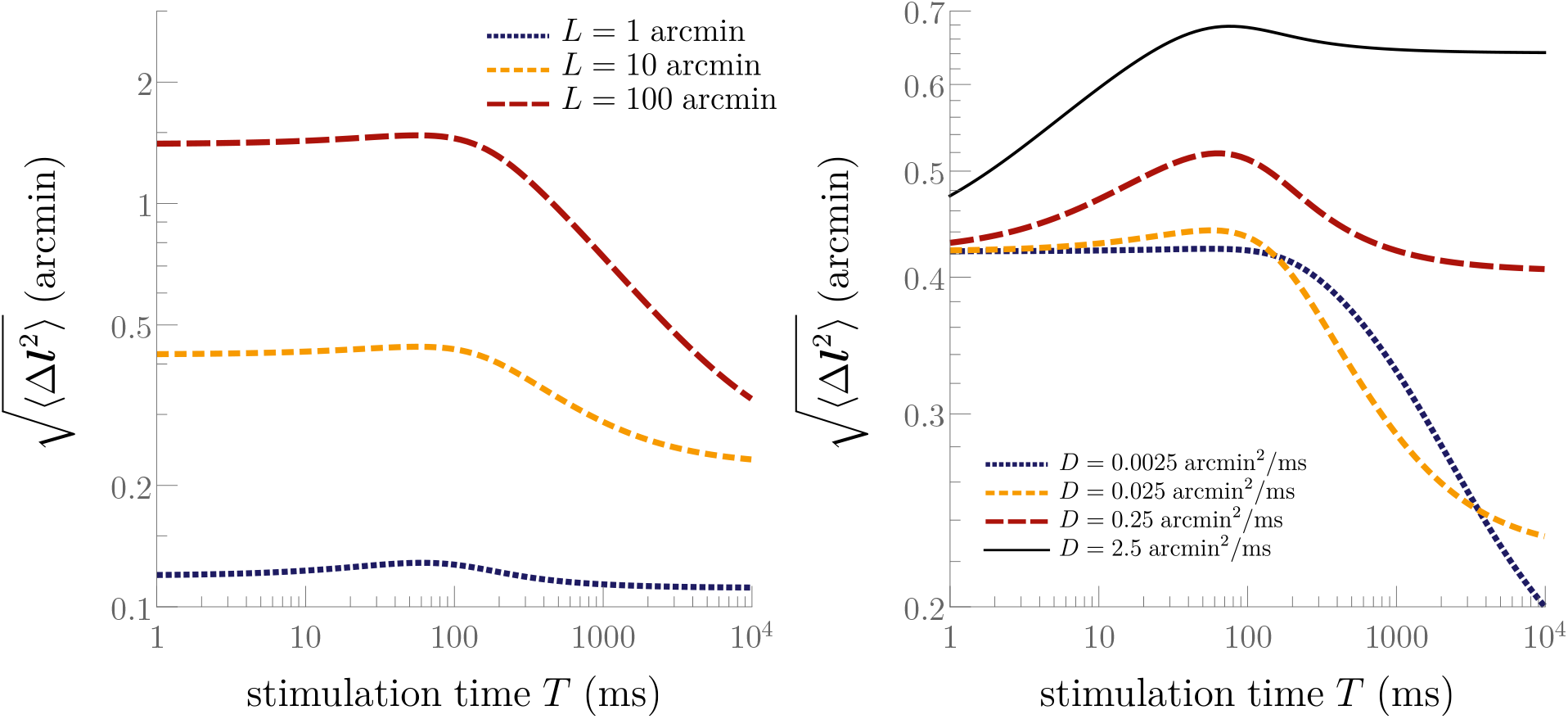
Left: Localization accuracy at the detection threshold (*z*_crit_ = 3) for diffusive eye movements with *D* = 0.025 arcmin^2^*/*ms corresponding to 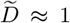 expressed as the standard deviation of the radially symmetric Gaussian distribution of likeliest locations *ϱ* (cf. Fig. 2). Even a briefly flashed stimulus once detected can be localized to a small fraction of its size. Larger stimuli exhibit a strong increase in accuracy with the onset of the detection plateau after about 100 ms. Right: Impact of the diffusion constant on localization of a stimulus of size 10’ at the corresponding detection threshold. Larger diffusion constants induce non-monotonic behavior of the localization accuracy. The black line represents eye movements optimized for detection. The yellow dashed lines in both panels are identical.

Importantly, even if the stimulus is shown only as a brief flash, if it is detected then it can also be localized to a high degree of accuracy. This justifies our high signal-to-noise ratio approximation which limits the applicability of our predicted localization accuracy to |Δ***l***| ≲ *β*. For a small stimulus this initial location uncertainty is a fixed fraction of the blur length 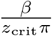, while for a large stimulus it increases with 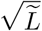, i.e. it is a decreasing fraction of the stimulus size. Moreover, if eye movements are tuned to support localization, the corresponding detectability of realistic stimuli always exhibits a plateau beyond the initial adaptation cycle. The majority of information for detection is hence obtained during the initial transient when the stimulus appears with *A*(*t*) as in eq. (1). During this period, eye movements have only a very limited effect on the acquired information so that detection is generally robust against fluctuations of eye movements.

The quantity depicted in Fig. 9 is the localization uncertainty which is not necessarily an easily accessible psychophysical observable. However, the Vernier task allows measuring localization accuracy projected onto an axis. While this introduces computational nuances between Vernier threshold and two-dimensional localization accuracy, in an isotropic setting we expect the length scale corresponding to the localization accuracy, i.e. 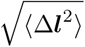 to qualitatively behave like a Vernier threshold. In that sense, Fig. 9 also describes the qualitative behavior of Vernier thresholds.

The experimental paradigm of measuring Vernier thresholds at fixed multiples of the detection threshold has been utilized across multiple studies, focussing on different stimulus parameters. Early studies found that Vernier thresholds continue to decrease for observation times longer than 150 ms [68–70]. This attracted attention in particular in comparison to detection tasks for which temporal summation past Bloch’s critical time was frequently reported as negligible [57, 71] sparking discussions that both tasks may be addressed by different mechanisms.

However, in [72], Vernier thresholds were measured at constant visibility, i.e. the contrast was adjusted relative to a line detection threshold as suggested in Fig. 9. This study found effectively no variation with stimulus duration of Vernier thresholds at fixed visibility. As the Vernier targets used were smaller than 1’ in the direction of offset and thus blur dominated, we expect the accuracy to be roughly constant in time as illustrated by the blue dotted line in Fig. 9. The matter is slightly complicated by the extent of Vernier targets perpendicular to the offset rendering them large in one dimension and small in the other. However, the integral in the direction perpendicular to the offset direction in the respective expressions for 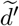 and 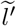 is identical and thus does not affect localization accuracy at constant detectability. Thus, our predictions are consistent with the observations in [72], which simplifies to another instance of detection and localization being effectively equivalent for small stimulus sizes. In order to resolve the effect of eye movements and adaptation, greater line widths are required.

Another general experimental observation is that localization accuracy increases roughly linearly with contrast in the vicinity of the contrast detection threshold [70, 73]. As our model is linear in contrast, this behavior is expected and supports our use of the approximation that transduction is close to linear near the detection threshold.

In [74], the authors investigate the effect of blur on Vernier acuity using lines with a Gaussian modulation in offset direction with variable standard deviation significantly larger than the intrinsic blur of the visual system. In the offset direction the stimulus is dominated by blur and can thus be considered point-like. Consequently, the standard deviation of the Gaussian target effectively replaces the blur parameter *β*. With the ratio 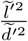 being constant for eye movements negligible compared to the blur length (cf. Fig. 7), the scaling of localization accuracy with *β* is given by eq. (23), i.e. *l*′ ∝ *β*^−1^. Accordingly, localization thresholds are expected to increase linearly with stimulus blur which is in excellent agreement with the experimental findings [74].

The right panel of Fig. 9 illustrates how the localization accuracy varies with eye movements. In the Bloch regime, any eye movement is detrimental but for sufficiently swift eye motion, localization suffers more than detection due to the B^∗^ regime (cf. Fig. 6). Accordingly, the location uncertainty increases with time for rapid eye motion. Beyond the Bloch regime, we observe the familiar decay in location uncertainty due to the plateau regime in the detection task. However, the larger the eye motion, the quicker the stimulus falls on a fresh part of retina, so the duration of the decay is reduced till it disappears entirely for 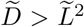. In the long time limit, the location uncertainty again approaches a plateau with a value that increases with 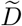 for 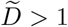 emphasizing that localization becomes relatively harder compared to detection when eye movements are too rapid.

Manipulating the extent of eye motion experimentally is not straightforward. However, Vernier acuity experiments have been conducted in stabilized conditions [68, 75] or with moderate added stimulus motion [76]. The general finding was that any kind of retinal motion has a remarkably small effect on Vernier acuity. Yet, all studies again feature Vernier bars small in the direction of offset for which any effect of eye movements is necessarily going to be small. As Fig. 9 illustrates, stabilization has a small effect even for larger stimuli compared to typical scale diffusive eye movements known exactly to the ideal observer. With additional stimulus motion, 100 ms stimulus duration Vernier thresholds are found mostly unaltered by imposed motion whereas for 200 ms presentations the data suggest a slight loss of acuity with larger velocities. Since motion also alters the detection threshold, which has not been accounted for in equating stimulus contrast across eye movement conditions, and since the stimuli employed have been small, these findings primarily relate to Fig. 5 which underlines little variation for diffusion constants smaller than optimal and a more substantial detriment for rapid eye movements. However, since stimulus movement may involve mechanisms different from fixation and stabilization may clash with the brain’s expectations of retinal motion, any experiment that artificially alters eye movements needs to be interpreted carefully. Alternatively, the acuity of different individuals with different eye movement characteristics may be compared. This is where the experimental paradigm of Vernier acuity at detection threshold becomes most powerful as it facilitates the comparison of performance across participants irrespective of their individual internal noise.

## V. CONCLUSION

In this article we demonstrated that the visual system’s tasks of detection and localization are closely related and can be analyzed within the same Bayesian framework for an ideal observer. A key result emerging from this analysis is that localization is equivalent to detection of the stimulus gradient, which establishes a fundamental link between both operations and facilitates the interpretation of all further results. We explored how the two tasks are affected by stimulus conditions (stimulus size and presentation time) and eye motion for a simple diffusive model. This allowed us to construct “scaling diagrams” for each task, which compactly summarize the emerging phenomenology, quantifying how performance in each task varies with stimulus conditions and eye motion. A rich scaling behavior arises through the interplay between the four relevant length scales of the system: the eye’s blur length, the distance traveled during one adaptation cycle, the distance traveled during stimulation, and the stimulus size. For stimuli smaller than the eye’s natural blur scale, performance in detection and localization tasks vary in almost identical ways with model parameters. Optimal eye motion for both tasks corresponds to moving roughly one blur length in one adaptation time. For larger stimuli, performance in detection and localization tasks vary differently with model parameters. Notably, information on localization always comes from the perimeter of a stimulus where gradients are largest. As a consequence, for localization of large stimuli, optimal motion still corresponds to moving roughly one blur length in one adaptation time. In contrast to that, for detection, at short presentation times the information is acquired from the whole stimulus area, being dominated by the temporal transients for turning the stimulus on and off. Optimal motion for detection corresponds to moving roughly the stimulus length in one adaptation time. Comparison with actual eye movements suggests that they are optimized for localization rather than detection. Our scaling predictions reproduce established theory such as Bloch’s, Ricco’s, and Piper’s law and additionally provide new explanations to experimental observations such as attributing fourth-root spatial summation to fixational eye movements. An insightful experimental protocol to emphasize the effect of fixational eye movements is to consider localization performance for stimuli presented with contrast adjusted to the detection threshold. Predicting the results of such a protocol requires understanding both detection and localization. As localizability and detectability are affected by stochastic fluctuations of the visual processing circuitry in the same way, the ratio of these performance measures depends predominantly on eye movements. Previous studies utilizing this paradigm were using small stimuli for which localization and detection become effectively equivalent so that any effect of eye movements is strongly suppressed. However, different eye movements interact differently with either task for stimuli large in comparison to visual system’s effective blur. Therefore, this experimental protocol in connection with large stimuli is a promising avenue to study the perceptual implications of fixational eye movements in combination with adaptation.

## ACKNOWLEDGMENTS

This work was supported by the UK Research And Innovation (UKRI) and the Wellcome Trust grant “A fresh look at visual sampling: How are fixational eye-movements optimised? [PhysFEM],” Grant No. EP/W023873/1. Generative AI (GPT-5.6 Sol) has been used to identify additional references on psychophysical experiments relevant to our theoretical predictions, and on localization methodology across domains.

## Appendix A: Bayesian Analysis of the Detection Task

Under the IO-assumptions we cast the detection task as a Bayesian inference. The IO knows that they are observing one of two conditions: Either the stimulus *s* is present, resulting in the noisy spatio-temporal excitation pattern *y*(***r***, *t*) ≡ *y*_on_(***r***, *t*) + *η*(***r***, *t*), or the stimulus is absent resulting in plain noise *y*(***r***, *t*) = *y*_off_ (***r***, *t*) + *η*(***r***, *t*) with *y*_off_ ≡ 0. The only stochastic component is the added Gaussian white noise. At this point it is helpful to think of individual receptors covering a square of length Δ*r* and discrete time steps Δ*t*. The IO can estimate the likelihood of any signal at discrete coordinates in space and time *y*(***r***_*i*_, *t*_*i*_) given the stimulus was on (resp. off) because the difference Δ*y* ≡ *y* − *y*_on_ is a Gaussian random variable with vanishing mean and variance 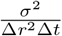. The noise variance needs to be scaled with the spatio-temporal “bin size” such that the average over that bin is characterized by the imposed variance *σ*^2^. That means, given the stimulus is “on” the probability to perceive *y*_*i*_ ≡ *y*(***r***_*i*_, *t*_*i*_) decays as

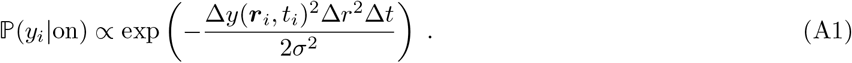

The realizations Δ*y*_*i*_ for different locations and times are independent. Thus, the likelihood of these realizations factorizes, i.e.,

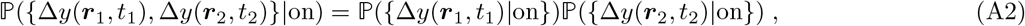

where the vertical bar indicates conditional probability. Hence, summing over all detectors and all times, the probability of obtaining the set of observations {*y*_*i*_} given the stimulus is “on” is:

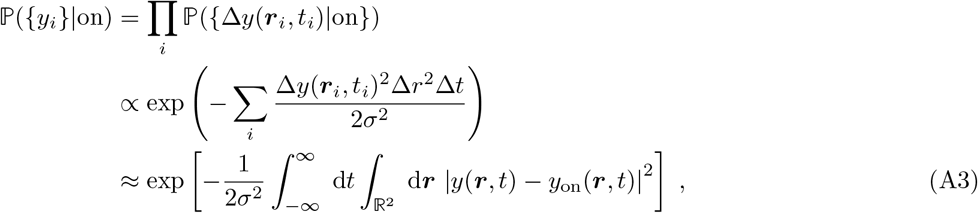

where the last approximation is made by taking the limits to continuous space and time Δ*r* → 0, Δ*t* → 0, so the sum becomes an integral. It is worth noting that we take this limit for computational ease going forward rather than physical motivation as the receptors on the retina are naturally discrete and time resolution of the neural circuit is likewise bounded.

Using Bayes’ theorem, we express the posterior probability of the stimulus being “on” given the set of signals of all receptors at all times {*y*_*i*_}

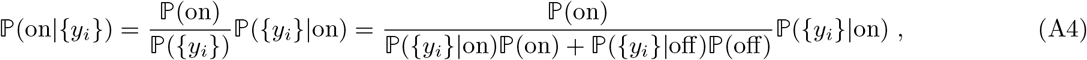

where ℙ(on) and ℙ(off) are the prior probabilities of the stimulus being on and off, that is, constants prescribed by the experimental setup. We assume these are equal, i.e. ℙ(on) = ℙ(off), but note that unequal prior probabilities are readily accounted for.

The IO simply picks the stimulus condition they deem more probable given a set of signals {*y*_*i*_}. Thus, the decision is made on the basis of the ratio

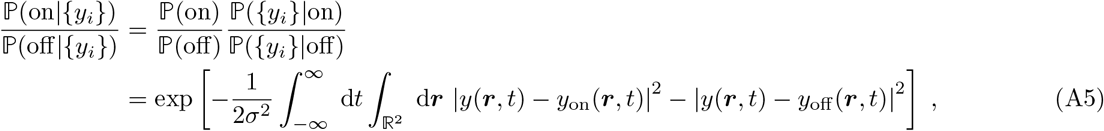

where we assumed ℙ(on) = ℙ(off). If this ratio is greater than 1, the IO will conclude the stimulus is on, and otherwise conclude the stimulus is off. Thus, the IO concludes the stimulus is on if:

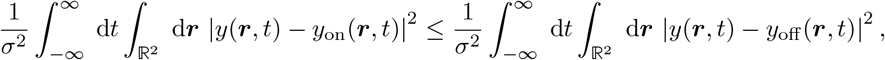

otherwise, the IO will conclude the stimulus is off. A correct decision corresponds to the IO deciding “on” if the stimulus was in fact on (or, conversely, deciding “off” if the stimulus was off). If the stimulus is on, then *y*(***r***, *t*) ≡ *y*_on_(***r***, *t*) + *η*(***r***, *t*) and so the IO correctly decides “on” if:

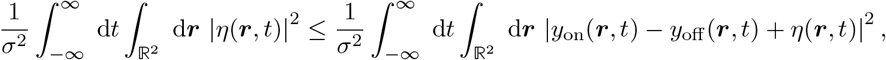

which simplifies to

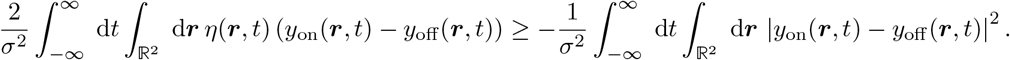

Thus, the decision is correct if *ξ >* −*d*^′2^, where

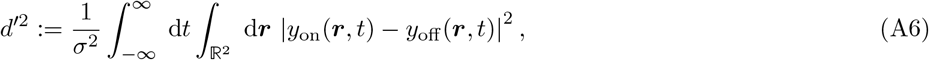

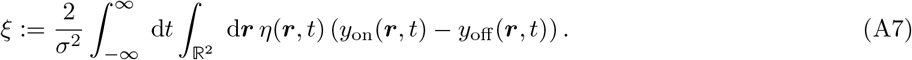

*d*^′2^ is purely deterministic, and quantifies the difference between the mean excitation patterns generated by the on and off conditions. *ξ* is a stochastic integral over scaled white noise which is distributed according to a normal distribution. Hence, *ξ* has mean zero and, making use of eq. (4), its variance can be calculated to equal 4*d*^′2^. Accordingly, we effectively draw a random variable *ξ* ~ *N*(0, 4*d*^′2^) and whenever *ξ >* − *d*^′2^, the IO makes correct decision. Integrating over all white noise realizations the probability of a correct decision depends exclusively on *d*^′2^:

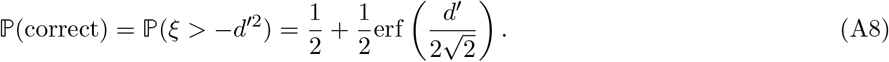

Although the last part of this analysis was performed assuming the stimulus to be on, the analysis with stimulus off gives the same result assuming equal prior probabilities of the stimulus condition.

## Appendix B: Distribution of the Position of Maximal Likelihood

In this Appendix we determine the position ***l*** of maximum likelihood for location of a stimulus, and quantify the error relative to the actual location ***l***_0_ of the stimulus. As given in eq. (15), for flat priors the posterior probability of ***l*** given the retinal response *y*_***l***_ (***r***, *t*)) is proportional to the likelihood ℙ (*y*_***l***_ (***r***, *t*)|***l***), i.e.

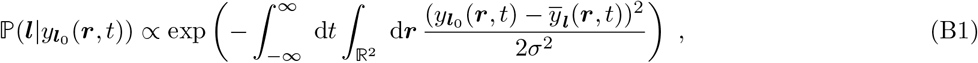

with 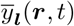 being the expected noise-free response for stimulus at position ***l***. We may write the likelihood in terms of the spatially Fourier transformed functions

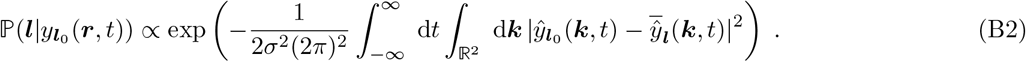

Using eq. (14), we write this in terms of the noise free response for a stimulus at the origin, 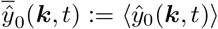, and noise 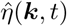. The corresponding log-likelihood is given up to an additive constant by

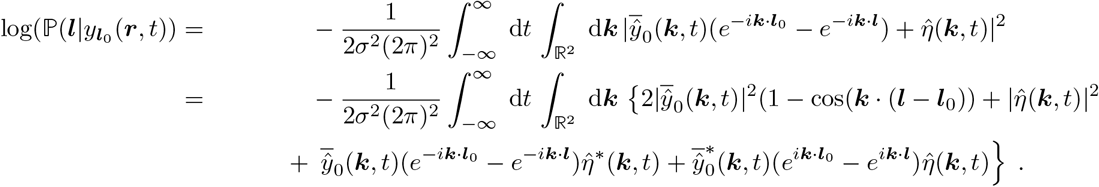

The location ***l*** that maximizes the likelihood, and therefore its logarithm, constitutes the best guess at the true stimulus location ***l***_0_ and satisfies

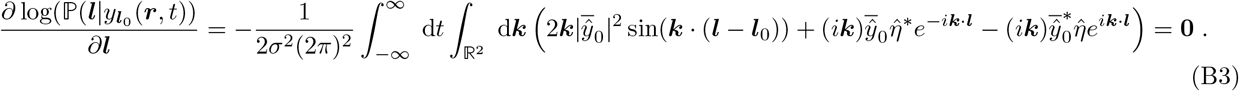

Once sufficient amounts of information have been acquired we can expect ***l*** to be reasonably close to the true location, such that in the relevant ***l***-region Δ***l*** := ***l*** − ***l***_0_ becomes small. The spectral weight function contains blur which naturally provides a large ***k***-cutoff. Thus, the relevant ***k***-range is finite so that eventually we can be confident in Δ***lk*** ≪ 1. Expanding the integrand of eq. (B3) to first order in Δ***lk*** and the noise we find

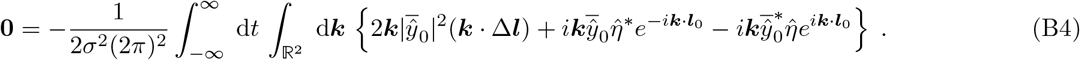

Isolating Δ***l***, we find

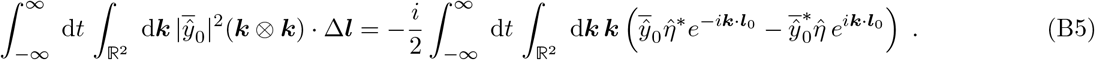

Compressing the operator on the l.h.s. into the symmetric tensor,

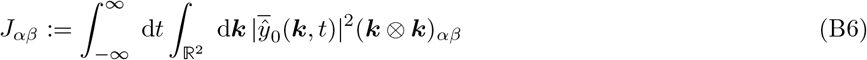

and the r.h.s. into a vector

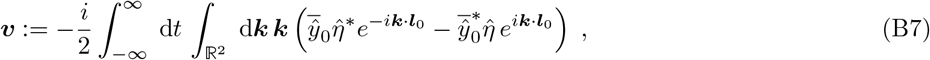

characterizing Δ***l*** formally simplifies to a matrix-vector multiplication

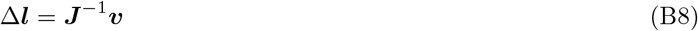

and average the result over the white noise realizations. Since the Fourier transformed white noise remains zero mean, then ⟨***v***⟩ = **0** in eq. (B8) which, assuming ***J*** is not singular, implies

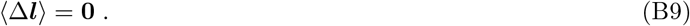

This is expected due to symmetry of the noise. However, utilizing the inherited properties of the Fourier transformed white noise

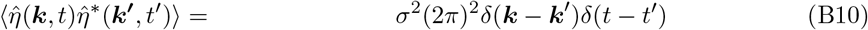

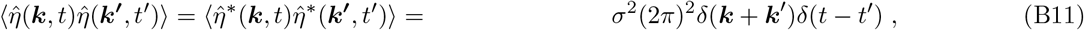

we may also compute the second moment of the distribution of Δ***l***. For the equivalent one-dimensional problem (Ω = ℝ) this is particularly simple and we readily find

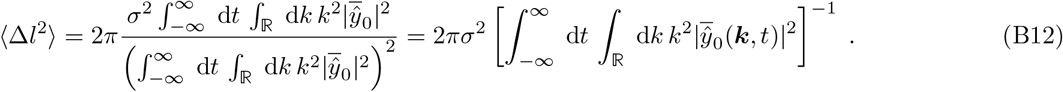

Naturally, we are more interested in two-dimensional retinas. In general, we need to compute

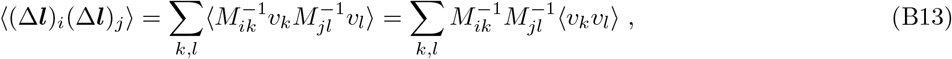

where only the vector ***v*** is affected by the noise fluctuations. Performing the average

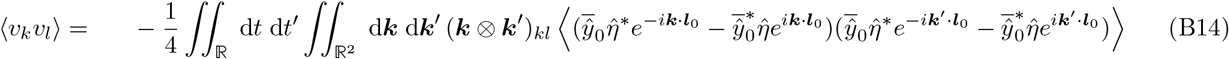

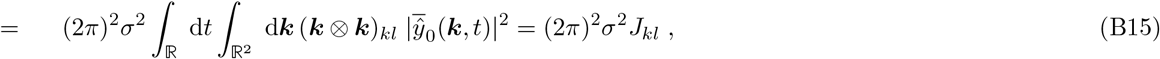

where the *δ*-distributions resulting from the noise variance enforce complex conjugation if necessary so that each of the four terms contributes equally. The noise average of the vector cancels one of the inverse matrices leaving us with the generalization of the one-dimensional result

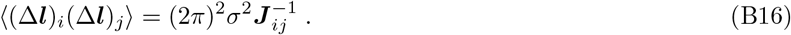

The degree to which a stimulus can be localized depends on the matrix ***J*** whose components are defined in eq. (B6). Expressing 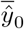 in terms of the baseline model gives:

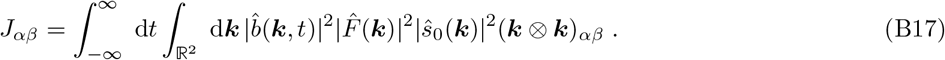

In the main text we absorb the prefactor into the definition of ***J*** so that eq. (B16) corresponds to the simpler form of eq. (16). These are the central results used in the main text.

## Appendix C: Model Specifications

### 1. Adaptation Kernel

The exact form of this kernel ultimately does not matter too much as long as it comprises the signature features: a short-time positive integration term and a long-time negative restoration term which integrate to zero (perfect adaptation), i.e. 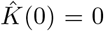. These conditions impose the general structure of a bandpass filter and beyond that we may pick an analytic form which keeps all expressions tractable. We choose a bi-exponential

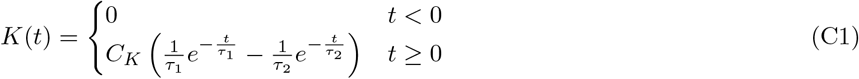

introducing the timescales 0 *< τ*_1_ *< τ*_2_ corresponding to integration and restoration, respectively. The constant *C*_*K*_ is arbitrary and simply scales the deterministic part of the signal so it effectively acts as an amplitude. Another reasonable normalization is 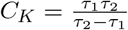 so that *K*(0) = 1 which is plotted below Fig. 10 to facilitate the comparison between different choices for the ratio 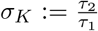. For the main part of this manuscript we set *C*_K_ = 1 to absorb this constant into the amplitude *A* of the stimulus. The corresponding power spectrum reads

**Figure 10.**
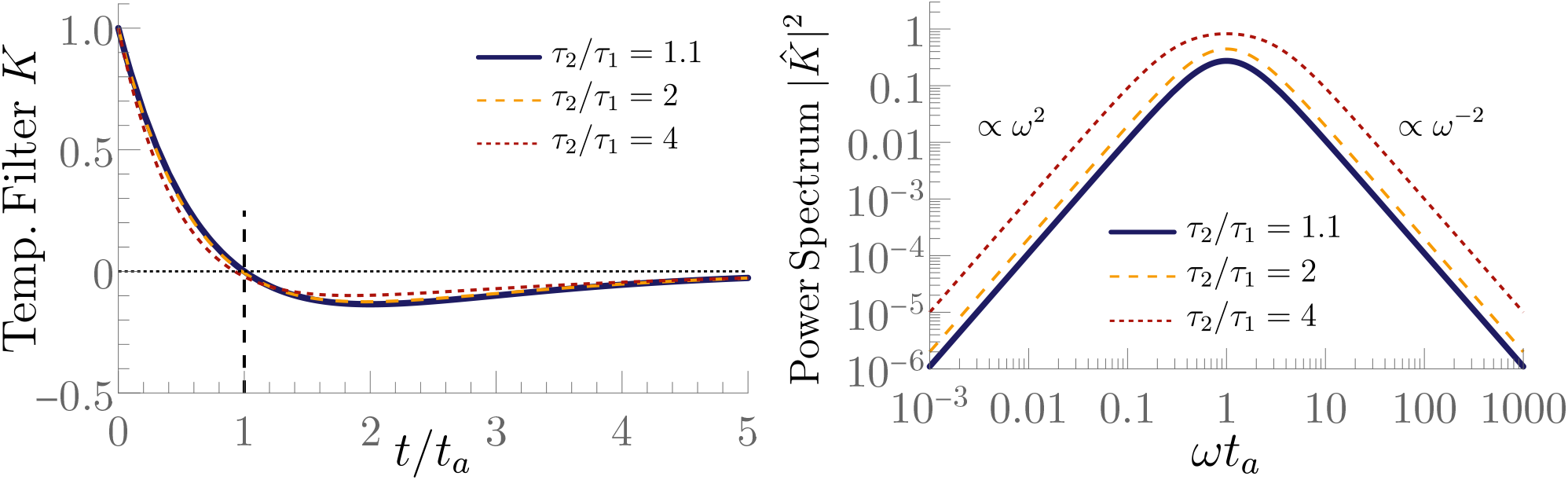
Time-dependence (left) and power spectrum (right) of the temporal filters used throughout this work. *K* is plotted with *τ*_2_ = 1.1 and *τ*_1_ = 1. eq. (C2) *K* has been normalized so that *K*(0) = 1 to facilitate visual comparison – the overall admitted intensity of both filter vastly differs with that normalization.

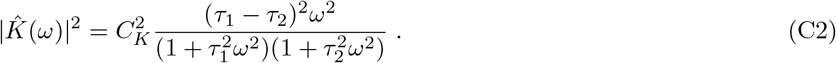

This family of bi-exponential kernels admits a characteristic time scale, the **adaptation time** which marks the root of *K* while *t* = 2*τ*_*a*_ is the locus of the minimum. Yet, this time depends rather intricately on both time scales – for computational ease we resort to the very similar geometric mean 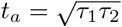. Throughout this study, we fix the ratio *σ*_*K*_ = 1.1 which next to *t*_*a*_ is the second parameter that is required to completely specify *K*. Yet, as is illustrated in Fig. 10, it has only a minor effect on the shape of the temporal filter and qualitatively the same power spectrum.

### 2. Stimulus

Stimulus and blurring function always appear as a product in Fourier space and depend exclusively on spatial frequencies if we associate the amplitude function with the eye movement. Thus, we can absorb the blurring function into a generalized stimulus. The most basic stimulus is a point which in connection with the blurring function yields a Gaussian in both real and Fourier space

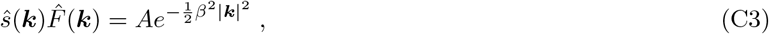

with *β*^2^ denoting the variance of the Gaussian blur. The spatial filter may again be arbitrarily normalized and the same applies for the stimulus. Together with the normalization constant of the temporal filter, we combine all these constant into the amplitude *A*. More intricately shaped stimuli *S* are integrals over point stimuli

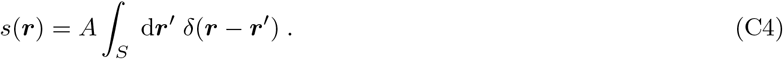

Consequently, the spatial Fourier transform is given by

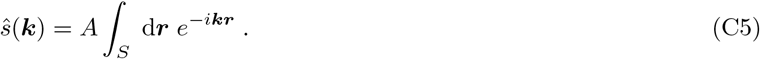

For the most part of the manuscript we consider a disk of radius *L* which has the power spectral density

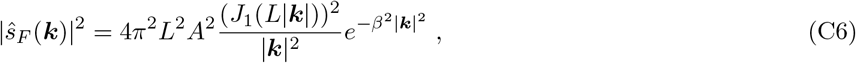

featuring the Bessel function *J*_1_. The appearance of the Bessel function renders completely analytical treatment cumbersome, the scaling results of section IV are obtained by using appropriate *k*-expansions. Much simpler is the study of rectangles of length *L* and width *W* as depicted in Fig. 4

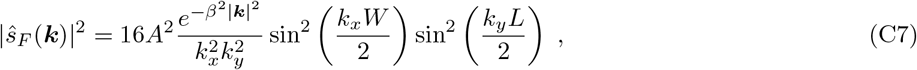

due to their separability. This becomes relevant when trying to connect our localization accuracy results to Vernier thresholds. As the Vernier task is only concerned with the localization accuracy projected onto one of the cardinal axes, the integrals in the perpendicular direction can be separated and coincide for detection and localization. In our proposed paradigm of studying localization at constant detectability that removes any effect of the perpendicular dimension as long as eye motion remains isotropic.

However, it swiftly gets more complicated when multiple rectangles are combined to form a tumbling E, a very common stimulus for discrimination tasks particularly for ophthalmological endeavors, despite its spectral complexity.

### 3. Eye Movements

We focus on the phenomenology induced by very simplistic building blocks indicating the suitability of certain patterns for specific tasks. These building blocks are **ballistic** eye motion, i.e. motion in a straight line characterized by a constant velocity ***v***.

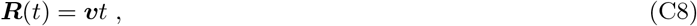

and **diffusive** eye motion, i.e. Brownian motion characterized by a diffusion constant *D* that controls the mean square displacement

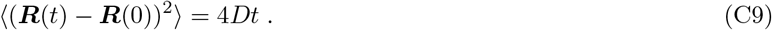

Even though realistic eye movements likely feature some degree of persistence, Brownian motion is the most studied instance of stochastic dynamics and thus an excellent benchmark model for more elaborate approaches. An important auxiliary result enabling the analytic treatment of the time integrals appearing in the expressions for 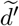 and 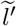 is

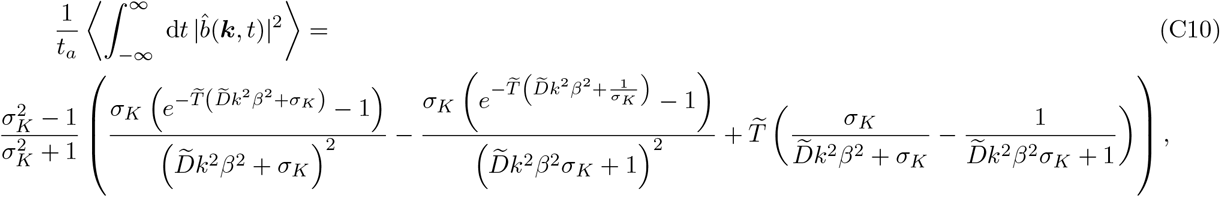

where ⟨·⟩ denotes an average over the ensemble of diffusive trajectories characterized by the diffusion constant *D*. This equation has been derived in [27] using slightly different terminology.

## Notes

### Competing Interest Statement

The authors have declared no competing interest.

### Summary of Updates

expanded the discussion of psychophysical experiments, added a discussion of the implications of the theoretical result beyond human vision, refined Figs. 2, 4 and 9, rephrased the abstract, improved general readability, removed typos

## References

[1] E. Pöppel, R. Held, and D. Frost, Residual visual function after brain wounds involving the central visual pathways in man, Nature 243, 295 (1973).

[2] S. L. Meeres and R. E. Graves, Localization of unseen visual stimuli by humans with normal vision, Neuropsychologia 28, 1231 (1990).

[3] P. Stoerig and A. Cowey, Blindsight in man and monkey., Brain: A Journal of Neurology 120, 535 (1997).

[4] D. M. Levi, S. A. Klein, and A. Aitsebaomo, Vernier acuity, crowding and cortical magnification, Vision Research 25, 963 (1985).

[5] H. R. Wilson, Model of peripheral and amblyopic hyperacuity, Vision Research 31, 967 (1991).

[6] H. Von Helmholtz, Helmholtz’s treatise on physiological optics, Vol. 3 (Optical Society of America, 1925).

[7] F. Ratliff and L. A. Riggs, Involuntary motions of the eye during monocular fixation., Journal of Experimental Psychology 40, 687 (1950).

[8] H. B. Barlow, Eye movements during fixation, The Journal of Physiology 116, 290 (1952).

[9] T. N. Cornsweet and H. D. Crane, Accurate two-dimensional eye tracker using first and fourth purkinje images, Journal of the Optical Society of America 63, 921 (1973).

[10] M. Rucci, E. Ahissar, D. C. Burr, I. Kagan, M. Poletti, and J. D. Victor, The visual system does not operate like a camera, Journal of Vision 25, 2 (2025).

[11] C. K. Sheehy, Q. Yang, D. W. Arathorn, P. Tiruveedhula, J. F. de Boer, and A. Roorda, High-speed, image-based eye tracking with a scanning laser ophthalmoscope, Biomedical Optics Express 3, 2611 (2012).

[12] W. M. Harmening, W. S. Tuten, A. Roorda, and L. C. Sincich, Mapping the perceptual grain of the human retina, Journal of Neuroscience 34, 5667 (2014).

[13] L. K. Young, T. J. Morris, C. D. Saunter, and H. E. Smithson, Compact, modular and in-plane aoslo for high-resolution retinal imaging, Biomedical Optics Express 9, 4275 (2018).

[14] M. Rucci and M. Poletti, Control and functions of fixational eye movements, Annual Review of Vision Science 1, 499 (2015).

[15] R. G. Alexander and S. Martinez-Conde, Fixational eye movements, in Eye Movement Research: An Introduction to its Scientific Foundations and Applications (Springer, 2019) pp. 73–115.

[16] S. Martinez-Conde, S. L. Macknik, and D. H. Hubel, The role of fixational eye movements in visual perception, Nature Reviews Neuroscience 5, 229 (2004).

[17] M. Ezenman, P. Hallett, and R. Frecker, Power spectra for ocular drift and tremor, Vision Research 25, 1635 (1985).

[18] N. Ben-Shushan, N. Shaham, M. Joshua, and Y. Burak, Fixational drift is driven by diffusive dynamics in central neural circuitry, Nature Communications 13, 1697 (2022).

[19] J. L. Reiniger, N. Domdei, F. G. Holz, and W. M. Harmening, Human gaze is systematically offset from the center of cone topography, Current Biology 31, 4188 (2021).

[20] R. J. Krauzlis, L. Goffart, and Z. M. Hafed, Neuronal control of fixation and fixational eye movements, Philosophical Transactions of the Royal Society B: Biological Sciences 372 (2017).

[21] A. B. Watson, Probability summation over time, Vision Research 19, 515 (1979).

[22] S. Laughlin, Coding efficiency and visual processing, in Vision: coding and efficiency (Cambridge University Press, 1990) pp. 26–31.

[23] B. H. Price and J. P. Gavornik, Efficient temporal coding in the early visual system: existing evidence and future directions, Frontiers in Computational Neuroscience 16, 929348 (2022).

[24] L. A. Riggs, F. Ratliff, J. C. Cornsweet, and T. N. Cornsweet, The disappearance of steadily fixated visual test objects, Journal of the Optical Society of America 43, 495 (1953).

[25] M. Rucci, Fixational eye movements, natural image statistics, and fine spatial vision, Network: Computation in Neural Systems 19, 253 (2008).

[26] J. Intoy and M. Rucci, Finely tuned eye movements enhance visual acuity, Nature Communications 11, 795 (2020).

[27] A. J. Houston, D. H. Brainard, H. E. Smithson, and D. J. Read, Information, movement, and adaptation in human vision, Physical Review Research 8, 023083 (2026).

[28] W. Bialek, Biophysics: Searching for Principles (Princeton University Press, 2012).

[29] A. B. Watson, K. Boff, L. Kaufman, and J. Thomas, Handbook of perception and human performance (1986).

[30] S. M. Smirnakis, M. J. Berry, D. K. Warland, W. Bialek, and M. Meister, Adaptation of retinal processing to image contrast and spatial scale, Nature 386, 69 (1997).

[31] H. De Vries, The quantum character of light and its bearing upon threshold of vision, the differential sensitivity and visual acuity of the eye, Physica 10, 553 (1943).

[32] A. Rose, The relative sensitivities of television pickup tubes, photographic film, and the human eye, Proceedings of the IRE 30, 293 (1942).

[33] A. Rose, The sensitivity performance of the human eye on an absolute scale, Journal of the Optical Society of America 38, 196 (1948).

[34] A. Kohn, Visual adaptation: physiology, mechanisms, and functional benefits, Journal of Neurophysiology 97, 3155 (2007).

[35] J. Benda, Neural adaptation, Current Biology 31, R110 (2021).

[36] C. A. Curcio, K. R. Sloan, R. E. Kalina, and A. E. Hendrickson, Human photoreceptor topography, Journal of Comparative Neurology 292, 497 (1990).

[37] F. A. Kingdom and N. Prins, Psychophysics: A practical introduction (Elsevier, 2016).

[38] N. V. S. Graham, Visual pattern analyzers (Oxford University Press, 1989).

[39] G. Westheimer, Visual acuity and hyperacuity., Investigative Ophthalmology & Visual Science 14 (1975).

[40] W. S. Geisler, Contributions of ideal observer theory to vision research, Vision Research 51, 771 (2011).

[41] D. M. Green and J. A. Swets, Signal Detection Theory and Psychophysics, Vol. 1 (Wiley New York, 1966).

[42] G. Westheimer, The spatial sense of the eye. proctor lecture., Investigative Ophthalmology & Visual Science 18, 893 (1979).

[43] M. Fahle and T. Poggio, Visual hyperacuity: spatiotemporal interpolation in human vision, Proceedings of the Royal Society of London. Series B. Biological Sciences 213, 451 (1981).

[44] W. S. Geisler, Physical limits of acuity and hyperacuity, Journal of the Optical Society of America A 1, 775 (1984).

[45] H. R. Wilson, Responses of spatial mechanisms can explain hyperacuity, Vision Research 26, 453 (1986).

[46] T.-A. E. Nghiem, J. L. Witten, O. Dufour, W. M. Harmening, and R. Azeredo da Silveira, Fixational eye movements as active sensation for high visual acuity, Proceedings of the National Academy of Sciences 122, e2416266122 (2025).

[47] V. Lukyanova, J. Ameln, J. L. Witten, A. Gutnikov, M. Freiberg, B. Sayim, and W. Harmening, Sub-cone visual acuity can be achieved with less than 1 arcmin retinal slip, Journal of Vision 26, 7 (2026).

[48] L. M. Wilcox, J. H. Elder, and R. F. Hess, The effects of blur and size on monocular and stereoscopic localization, Vision Research 40, 3575 (2000).

[49] W. S. Geisler and K. D. Davila, Ideal discriminators in spatial vision: two-point stimuli, Journal of the Optical Society of America A 2, 1483 (1985).

[50] D. Robinson and P. Milanfar, Fundamental performance limits in image registration, IEEE Transactions on Image Processing 13, 1185 (2004).

[51] X. Kuang, M. Poletti, J. D. Victor, and M. Rucci, Temporal encoding of spatial information during active visual fixation, Current Biology 22, 510 (2012).

[52] R. Engbert, K. Mergenthaler, P. Sinn, and A. Pikovsky, An integrated model of fixational eye movements and microsaccades, Proceedings of the National Academy of Sciences 108, E765 (2011).

[53] J. A. Roberts, G. Wallis, and M. Breakspear, Fixational eye movements during viewing of dynamic natural scenes, Frontiers in Psychology 4, 797 (2013).

[54] A. Gorea, A refresher of the original bloch’s law paper (bloch, july 1885), i-Perception 6, 2041669515593043 (2015).

[55] O. Lévêque, C. Kulcsár, L. Cognet, and F. Goudail, On the equivalence of binary phase masks optimized for localization or detection in extended depth-of-field localization microscopy, Journal of the Optical Society of America A 40, 1753 (2023).

[56] D. Lang and D. W. Hogg, Principled point-source detection in collections of astronomical images, The Astronomical Journal 171, 231 (2026).

[57] H. Barlow, Temporal and spatial summation in human vision at different background intensities, The Journal of Physiology 141, 337 (1958).

[58] V. Glezer, The receptive fields of the retina, Vision Research 5, 497 (1965).

[59] T. Inui, O. Mimura, and K. Kani, Retinal sensitivity and spatial summation in the foveal and parafoveal regions, J. Opt. Soc. Am. 71, 151 (1981).

[60] G. W. Davila KD, The relative contributions of pre-neural and neural factors to areal summation in the fovea, Journal of Vision 31 (1991).

[61] S. K. Khuu and M. Kalloniatis, Spatial summation across the central visual field: Implications for visual field testing, Journal of Vision 15, 6 (2015).

[62] W. S. Tuten, R. F. Cooper, P. Tiruveedhula, A. Dubra, A. Roorda, N. P. Cottaris, D. H. Brainard, and J. I. W. Morgan, Spatial summation in the human fovea: Do normal optical aberrations and fixational eye movements have an effect?, Journal of Vision 18 (2018).

[63] H. Piper, Uber die abhangigkeit des reizwertes leuchtender objekte von ihrer flachen-bezsw. winkelgrosse, Zeitschrift fur Psychologie und Physiologie der Sinnesorgane 32, 98 (1903).

[64] K. D. Davila and W. S. Geisler, The relative contributions of pre-neural and neural factors to areal summation in the fovea, Vision Research 31, 1369 (1991).

[65] Y. Bonneh and D. Sagi, Contrast integration across space, Vision Research 39, 2597 (1999).

[66] T. S. Meese, R. F. Hess, and C. B. Williams, Size matters, but not for everyone: Individual differences for contrast discrimination, Journal of Vision 5, 2 (2005).

[67] A. S. Baldwin and T. S. Meese, Summation of contrast across the visual field: a common “fourth root” rule holds from the fovea to the periphery, Vision Research 243, 108784 (2026).

[68] U. T. Keesey, Effects of involuntary eye movements on visual acuity, Journal of the Optical Society of America 50, 769 (1960).

[69] W. S. Baron and G. Westheimer, Visual acuity as a function of exposure duration, Journal of the Optical Society of America 63, 212 (1973).

[70] S. J. Waugh and D. M. Levi, Visibility, timing and vernier acuity, Vision Research 33, 505 (1993).

[71] R. M. Saunders, The critical duration of temporal summation in the human central fovea, Vision Research 15, 699 (1975).

[72] S. J. Waugh and D. M. Levi, Spatial scale of visual analysis for vernier acuity does not vary over time, Vision Research 40, 163 (2000).

[73] C. A. Burbeck, Exposure-duration effects in localization judgments, Journal of the Optical Society of America A 3, 1983 (1986).

[74] J. Krauskopf and B. Farell, Vernier acuity: effects of chromatic content, blur and contrast, Vision Research 31, 735 (1991).

[75] D. H. Fender and P. Nye, The effects of retinal image motion in a simple pattern recognition task, Kybernetik 1, 192 (1962).

[76] G. Westheimer and S. P. McKee, Visual acuity in the presence of retinal-image motion, Journal of the Optical Society of America 65, 847 (1975).

